# Early neuronal reprogramming and cell cycle reentry shape Alzheimer’s disease progression

**DOI:** 10.1101/2025.06.04.653670

**Authors:** Roi Meir, Gali Schwartz, Miriam Adam, Adi Avni Lapidot, Anael Cain, Gilad Sahar Green, Vilas Menon, David A. Bennett, Philip L. De Jager, Naomi Habib

## Abstract

Alzheimer’s disease (AD) is a progressive neurodegenerative disorder characterized by hallmark pathologies, synaptic dysfunction, neuronal loss, gliosis and cognitive decline-dementia. Recent large-scale cell atlases of human AD brains exposed vulnerability of specific neuronal subtypes and highlighted early, coordinated glial responses, suggesting glial involvement in disease progression. However, the timing and nature of neuronal changes, differences between neuronal subtypes, and their coordination with glia remain unclear. Here, we used non-negative matrix factorization to identify co-expression gene programs in single-nucleus RNA profiles from the prefrontal cortex of 437 samples from donors whose clinical symptoms ranged between no cognitive impairment and AD dementia. This approach identified early coordinated transcriptional changes across all neuronal subtypes, preceding clinical symptoms of cognitive decline, and validated in independent snRNA-seq, proteomics, and ELISA datasets. We found neurons in AD undergo rapid modulation of synaptic genes, accompanied by convergence of neurons into two distinct programs: An oxidative stress and apoptosis program abundant in vulnerable neuronal subtypes, and a DNA damage and cell-cycle reentry program associated with resilient subtypes. Moreover, neuronal reprogramming was closely tied to glial responses, and diverged between AD to non-AD brain aging, suggesting neuro-glial coordinated reprogramming shapes the AD cascade and influences disease outcomes.

## Introduction

Alzheimer’s Disease (AD) is a progressive neurodegenerative disease characterized, beyond neuronal loss and synaptic dysfunction, by the accumulation of hallmark pathologies and gliosis, which emerge at the pre-clinical stage of AD prior to the clinical symptoms of cognitive-decline and dementia^1^. Large-scale profiling of post-mortem aging brains with single-cell resolution by single nucleus RNA-seq (snRNA-seq)^2,3^, uncovered rapid reprogramming of the transcriptional state of glial cell types along the progression of the disease^4–10^. Moreover, these cascades of cellular changes are coordinated across cell types, forming cellular communities of correlated cell subpopulations that characterize disease progression^5,6^. Furthermore, divergent glial responses were recently identified to distinguish between the progression of AD (prAD) and of alternative brain aging (ABA), linking the glial states to disease outcomes^6^.

Despite the link between synaptic dysfunction and neuronal loss to the cognitive symptoms of AD, large-scale omics studies have highlighted glial dynamics, leaving the timing and modulation of neuronal changes—particularly across neuronal subtypes largely unknown. Emerging data suggest selective vulnerability of specific subtype of neurons, showing a higher rate of loss of Somatostatin (SST) inhibitory neurons^5–8^ and a lower rate of loss of Parvalbumin (PV) inhibitory and L2-3 IT excitatory neurons^7^. While some molecular and physiological differences between these subtypes are known^11^, the drivers of selective vulnerability remain unclear. Identifying neuronal gene expression and mapping their dynamics along AD progression across neuronal subtypes may address these critical open questions and provide insights into the mechanisms driving neurodegeneration.

We hypothesized that standard sc/snRNA-seq analysis hinders the discovery of transcriptional changes in AD, as these approaches assign cells to discrete clusters based on the strongest axes of variation, such as stable neuronal subtypes^12^, potentially overlooking the complexity of cells executing multiple parallel functions and missing subtle, shared responses across neurons.

To capture the complex multifaceted cellular responses along the AD cascade, specifically in neurons, we applied a continuous approach, using non-negative matrix factorization (NMF) to model cells as a mixture of co-expressed gene programs in a large snRNA-seq atlas from 437 aging brains. We identified neuronal programs shared across neuronal classes that change along the AD cascade, and are validated at the protein level. These changes emerge at early stages of AD, and are highly coordinated with glial responses. Notably, we uncovered two neuronal programs increasing in AD that are mutually exclusive at the single cell level, reflecting different neuronal fates: one linked to stress responses and neuronal apoptosis, and the other to DNA damage and cell-cycle reentry and arrest. We linked the cell-cycle reentry program to the resilient neuronal subtypes and the stress/apoptotic program to the vulnerable subtypes. Our work points to a coordinated neuronal-glial reprograming which drives AD progression, aligning specific functions along the disease timeline which distinguish AD from brain aging, and pointing to a critical stage of the disease where glial responses converge with sharp loss of synaptic proteins.

## Results

### Continuous modelling capture programs shared across neuronal subtypes in the aging human cortex

To capture the diversity of cellular functions and the coordinated modulations and dynamics of neuronal and glial cells in AD and aging, we generated an atlas of cell-type specific co-expression gene programs in the human dorsolateral prefrontal cortex (DLPFC). We applied an NMF-based fast-topic algorithm^13^ to 1.58 million single nucleus RNA profiles from 437 aging individuals covering a wide range of healthy and pathological aging, including all stages of AD^6^ (from the ROS/MAP cohort^14,15^, **Methods**, **Fig. 1a, Supplementary Table 1**). In a continuous modelling approach, cells are represented as a combination of co-expressed gene programs, rather than assigning each cell to a single cellular subset (cluster), thereby enabling cells to be similar along some axes yet distinct along others, and capture both subtype identity and transient cellular responses within neuronal and glial cells (**Fig. 1b**).

**Figure 1.**
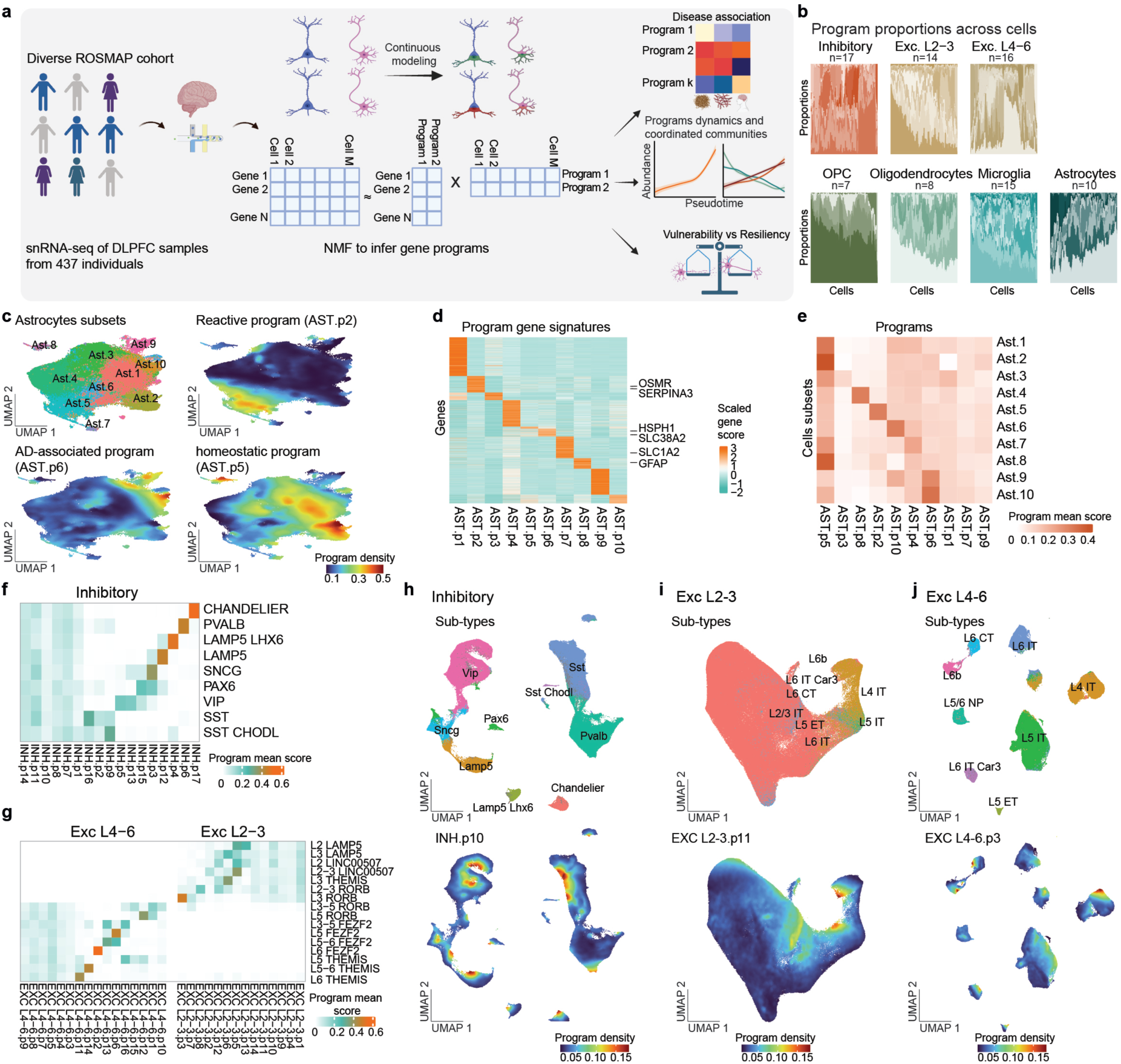
Cell-type specific co-expression atlas of the aging DLPFC capture shared and subtype-specific neuronal programs. **a**. Schematic overview of the experimental design and the analytical workflow enabling the construction of cell-type specific co-expression atlas of the human aging DLPFC. **b.** Expression programs capture cellular diversity. Proportions of programs (colour shades) per cell (columns) within each cell type (over 5000 random cells per type). **c**. Astrocyte expression programs capture distinct subsets and align with clusters. 2D UMAP embedding of (n=228,925) snRNA-seq profiles of astrocytes coloured by: previously defined astrocytes clusters^6^ (subsets, top left) and by locally smoothed density scores of representative programs - reactive AST.p2 (Ast.5^6^-like), disease associated stress response AST.p6 (Ast.10^6^-like) and homeostatic AST.p5 (Ast.1-2^6^-like). **d.** Unique expression signatures of astrocyte programs. Gene score (F, **Methods**) for top differentially expressed genes (rows, scaled) across programs. **e.** Astrocyte programs align with clusters. Averaged program scores in each cell subset (clusters from ^6^). **f-j.** Neuronal programs capture subtype-specific and shared programs: **f,g.** Unique expression signatures of inhibitory (f) and excitatory (g) neuronal programs (as in d). **h,i,j.** 2D UMAP embedding of snRNA-seq profiles of inhibitory (h, n=257,929), excitatory upper layers (i, n=378,546) and excitatory lower layers (j, n=268,854), coloured by reference neuronal subtypes (published clusters^6^ annotated by Allan Brain Map^10^, top) and by a representative shared program (bottom).

Each *program* is defined by assigned score per gene and by a unique *gene signature* (differentially expressed genes, **Methods, Supplementary Table 2**), and each cell is assigned a score per program (**Fig. 1b**). We show that our atlas is robust to the number of programs (**Extended Data Fig. 1a,b**) and to sub-sampling of cells or neuronal subtypes (**Extended Data Fig. 1c,d)**, and that all programs are shared across multiple individuals (**Extended Data Fig. 1e, Methods**). We defined between 5 to 17 gene programs per cell type, capturing programs shared across all cells as well as programs that are unique to a subset of cells (**Fig. 1b,c Extended Data Fig. 2**).

In glial cells, our programs-based annotations for astrocytes (AST, 10 programs), microglia (MIC, 15 programs), oligodendrocytes (OLI, 8 programs) and oligodendrocytes precursor cells (OPC, 7 programs) largely recapitulated previously defined cellular states defined by clustering^6^ and further refined annotations by untangling overlapping biological functions (**Fig. 1c-e, Extended Data Fig. 2**). We identified homeostatic programs (*e.g.* AST.p5 and MIC.p9 programs matching previously defined clusters Ast.1-Ast.2^6^ and Mic.2-4^6^, respectively), multiple reactive programs (*e.g.* reactive programs AST.p2, MIC.p5 and MIC.p15 matching clusters Ast.5, Mic.6-7^6^, respectively), interferon response programs (*e.g.* AST.p4, matching cluster Ast.7^6^), and AD-associated stress responding programs (*e.g.* AST.p6 and OLI.p8 matching clusters Ast.9-10^6^ and Oli.7^6^, respectively). Our programs further refined existing cluster assignment, distinguishing between overlapping biological functions. For example, the disease-associated microglia clusters (Mic.12 and Mic.13^6^) were captured by two partially overlapping programs – a lipid/foam associated program MIC.p11 and an inflammatory/immune program MIC.p8 (expressing multiple DAM markers^16^, **Extended Data Fig. 2a-c**).

For neuronal cells, the flexible modeling of inhibitory (INH, 17 programs), excitatory upper layers (EXC.L2-3, 14 programs) and excitatory lower layers (EXC.L4-6, 16 programs) neurons, uncovered both *neuronal subtype-specific programs* as well as novel *shared programs* reflecting broader responses across multiple neuronal sub-types (**Fig. 1b,f-j Extended Data Fig. 3**). Subtype-specific programs are specifically expressed within one/few defined neuronal subtype annotations^12^ and include known subtype markers, such as inhibitory subtypes SST captured in INH.p16 and PVALB captured in INH.p6, and similarly for excitatory laminar subpopulations. While, the shared programs are expressed across all neuronal subtypes within a neuronal class, such as INH.p10, INH.p7, EXC.L2-3.p11 and EXC.L4-6.p3 (**Fig. 1f-j Extended Data Fig. 3a,b**). To validate these shared programs, we re-fitted programs within individual inhibitory subtypes (*e.g.* SST and PV), which identified similar programs independently within each subtype, confirming our observations of shared neuronal reprogramming (**Extended Data Fig. 3c-e**).

### Coordinated neuronal reprogramming emerges early within the AD cascade

To identify programs associated with AD for each cell type, we regressed a participant’s mean program score against their AD pathology loads - cortical amyloid β (Aβ) load and cortical paired helical filaments tau (tau) density – as well as the rate of cognitive decline, while controlling for sex, age at death, and post-mortem interval (PMI) (FDR<0.05, **Fig. 2a, Methods, Supplementary Table 3**). We uncovered multiple glial programs associated with AD traits consistent with previous findings^4–6,8^, positive for disease-associated programs AST.p6, MIC.p8, MIC.p11 and OLI.p8, and negative for homeostatic programs AST.p5, MIC.p9, and OPC.p5. Additionally, multiple neuronal subtypes-specific programs as well as shared neuronal programs were significantly associated with AD traits (**Fig. 2a**). The associations of neuronal subtype-specific programs with AD traits aligned with prior reports of higher vulnerability or resilience of specific subtypes^5–7^. These included negative association of programs reflecting the vulnerable subpopulation of SST (INH.p16), and a positive association of the resilient INH.p6 subpopulation of PV and EXC.L2-3.p7 upper layers L2-3 IT neurons (**Fig. 2a**). Interestingly, we uncovered within each neuronal class, both positively and negatively AD-associated shared neuronal programs, such as the upper neuronal layers program EXC.L4-6.p4, lower layers EXC.L2-3.p9, and inhibitory neurons INH.p1, all negatively associated with Aβ and cognitive decline (**Fig. 2a**).

**Figure 2.**
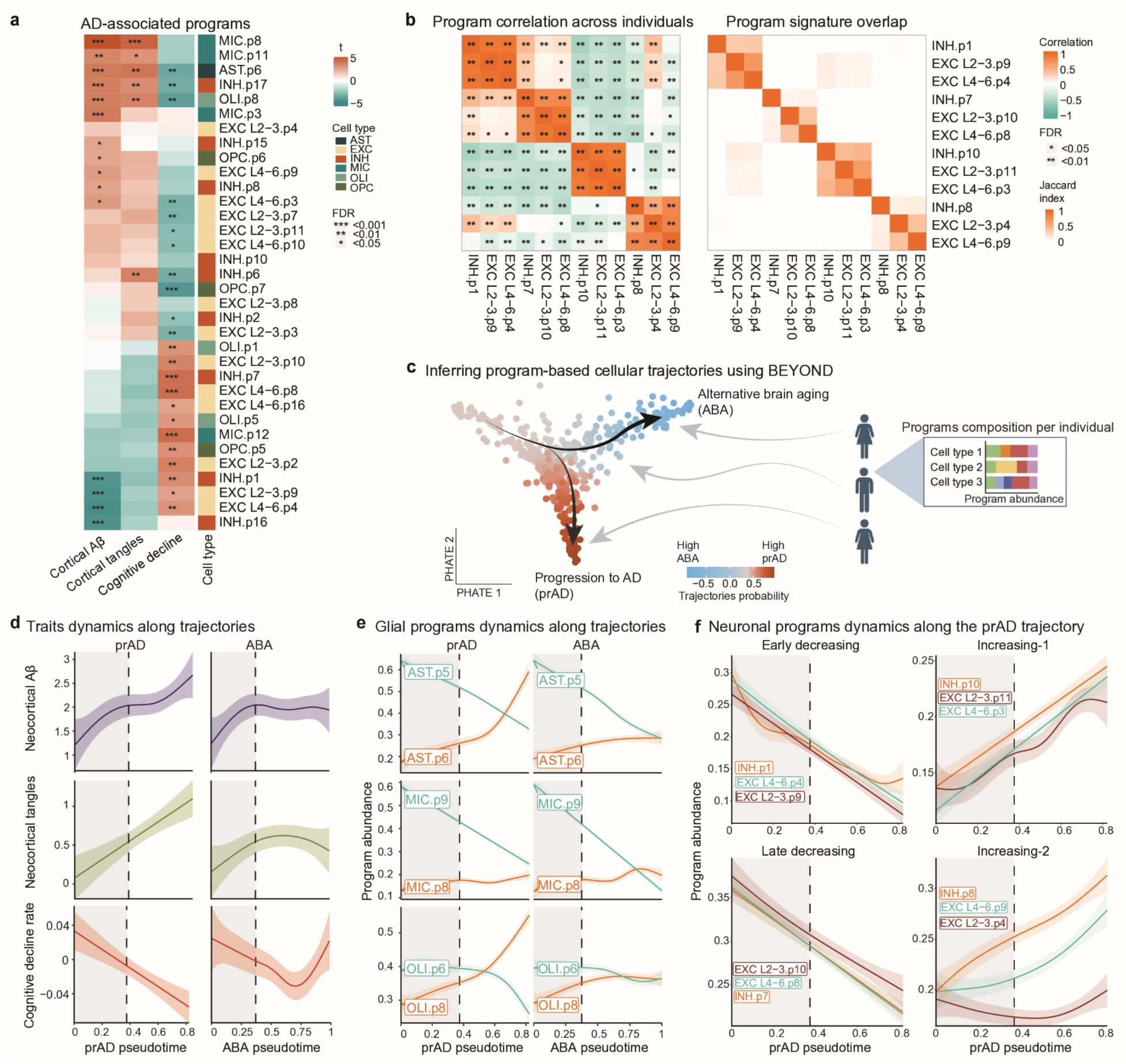
AD-associated neuronal programs are coordinated across neuronal classes. **a.** Association of programs to AD traits. Association of averaged program score across cells per individual and AD traits: cortical Aβ load, cortical tau load and rate of cognitive decline (linear regression controlled for confounders, **Methods**). * = FDR<0.05, ** = FDR<0.01, *** = FDR<0.001; **b.** Similarities of coordinated neuronal programs across classes. Left: Spearman correlation of averaged program scores across participants. Right: Jaccard similarity between program signatures of neuronal programs. **d.** Manifold of cellular programs captured two trajectories leading to prAD and ABA. PHATE^36^ embedding of each participant (dots) in the cellular landscape manifold captured by BEYOND^6^ algorithm based on their programs compositions (**Methods**). Dots are coloured by participants probability to be in the prAD and ABA trajectory. **e-f**. Dynamically regulated glial and neuronal programs along the prAD and ABA trajectories. Inferred dynamics of averaged program scores per individual over the pseudotime. Dashed line represents the split point between trajectories. Confidence intervals of standard error are displayed as lighter color. **e**. Decreasing homeostatic and increasing disease-associated programs in astrocytes (top), microglia (middle) and oligodendrocytes (bottom). **f**. Four groups of corresponding neuronal programs across classes: early-decreasing (top left), late-decreasing (bottom left), increasing-1 (top right) and increasing-2 (bottom right).

We further showed that these disease-associated neuronal programs are not only shared across neuronal subtypes, but rather are highly coordinated between the excitatory and inhibitory neuronal classes. Comparison across neuronal classes revealed four groups of *corresponding disease-associated neuronal programs* that spanned all three neuronal classes (**Fig. 2b**). These corresponding programs exhibited strong inter-individual concordance of their abundance (**Methods, Fig. 2b**) as well as overlapping gene signatures (especially among the excitatory programs, **Fig. 2b**), indicating a shared coordinated neuronal response in disease across neuronal classes.

To align the neuronal reprograming with disease stages, we inferred the dynamics of programs along the continuous trajectory of disease progression, revealing vast coordinated neuronal reprograming early in the disease cascade. To this end, we applied the BEYOND^6^ framework, which aligns individuals along pseudo-temporal trajectories of cellular change (**Fig. 2c, Methods, Supplementary Table 4**). As each post-mortem sample captures a snapshot of the cells at the time of death, it is assigned to a pseudotime along a trajectory, enabling inference of cellular and pathology dynamics by integrating data across individuals. BEYOND previously untangled the trajectories of AD progression and alternative brain aging when trained on snRNA-seq cluster abundance^6^. Here, we applied BEYOND on the abundance of neuronal and glial gene programs in 437 individuals (mean program score across all cells, **Methods**), identifying two distinct trajectories that recapitulated the published cluster-based trajectories^6^: one captured the gradual progression from healthy state to advanced AD (prAD) and another captured Alternative Brain Aging (ABA) (**Fig. 2c, Extended Data Fig. 4a,b**). Fitting clinicopathological traits dynamics along these two trajectories (**Method**), we indeed observed an increase in cortical Aβ and tau loads and acceleration of cognitive decline rate along the prAD trajectory, whereas the ABA trajectory showed lower levels of Aβ, limited tau load and variable cognitive decline rate (**Fig. 2d, Extended Data Fig. 4c)**.

Fitting the dynamics of gene programs revealed that the disease-associated glial and neuronal programs gradually changed in abundance specifically along the prAD trajectory, but not in the ABA (**Fig. 2e,f**). The increase of the glial disease-associated programs and the rapid decrease of the homeostatic programs, matched previous reports^6–8,10^ (**Fig. 2e, Extended Data Fig. 4d**) and recapitulated similar dynamics along published disease trajectories (**Extended Data Fig.4d**). All the *corresponding disease-associated neuronal programs* had similar dynamics along the prAD trajectory (**Fig. 2f, Extended Data Fig. 4e**). These *corresponding programs* were divided into four groups: an *early-decreasing* group of *programs* negatively associated with Aβ and with cognitive decline (INH.p1, EXC.L2-3.p9 and EXC.L4-6.p4), a *late-decreasing group* associated with cognitive decline (INH.p7, EXC.L2-3.p10 and EXC.L4-6.p8), an *increasing-1* group of programs associated with cognitive decline (INH.p10, EXC.L2-3.p11 and EXC.L4-6.p3) and *increasing-2* group positively associated with Aβ load (INH.p8, EXC.L2-3.p4 and EXC.L4-6.p9). Of note, INH.p10 and EXC.L2-3.p4 were only marginally associated with AD traits.

### Early synaptic dysregulation and metabolic shifts in neurons accompanying stress response along AD progression

To decipher the biological underpinnings of the neuronal programs, we performed pathway enrichment analysis on the signatures of each program (FDR<0.05, **Methods, Supplementary Table 2**), exposing a shared modulation across neuronal classes in disease in each of the four *corresponding disease-associated neuronal programs* (**Fig. 3a,b, Extended Data Fig. 5a)**. The *corresponding early-decreasing* programs were enriched in nuclear-encoded mitochondrial ribosomal genes, (previously linked to AD^7,8^, **Fig. 3c, Extended Data Fig. 5b**), antisense genes (*e.g*. *SNAP25-AS1*) and partially with mitophagy genes (*e.g. PINK1,TOMM5,TOMM7)* **(Fig. 3a,b, Extended Data Fig. 5c,d**), which may reflect early mitochondrial dysfunction or a metabolic shift and was shown to trigger the mitochondrial unfolded protein response (mt-UPR)^17^. The *corresponding late-decreasing* neuronal programs were enriched with postsynaptic specialization, synaptic structure pathways (*e.g. DLGAP4, GRIN2D, SHANK1*) and nuclear pore complex genes (**Fig. 3c, Extended Data Fig. 5e,f**). The *increasing-1* programs, were enriched with stress-associated responses, specifically metal ions and oxidative stress (*e.g. PRDX1, PRDX2*), mitochondrial genes, and the apoptotic processes (*e.g. BNIP3, HIPK2*), as well as pre-synaptic processes - mainly vesicle release (*e.g. VAMP2, SNAP25, CPLX1, CPLX2*), (**Fig. 3b,c, Extended Data Fig. 5g, Extended Data Fig. 6**). Finally, the corresponding increasing-2 neuronal programs were enriched in chromatin remodeling genes (*e.g. SMARCA1, SMARCAD1, CHD1*), RNA-splicing (*e.g. RNPC3, SRPK1, SRSF1*), heat shock response (*e.g. HSPH1, HSPA12A, HSPA4*), DNA damage repair response (*e.g. ERCC5, ATM, POLK*), as well as cell-cycle pathways, specifically M phase (*e.g. CENPC, ANAPC16, CEP290*), (**Fig. 3b,c, Extended Data Fig. 7a-d**). At the gene level, the similarity across neuronal classes varies between the programs and the classes, with greater similarity within the two excitatory neurons compared to the inhibitory (**Fig. 2b and Extended Data Fig. 5a**).

**Figure 3.**
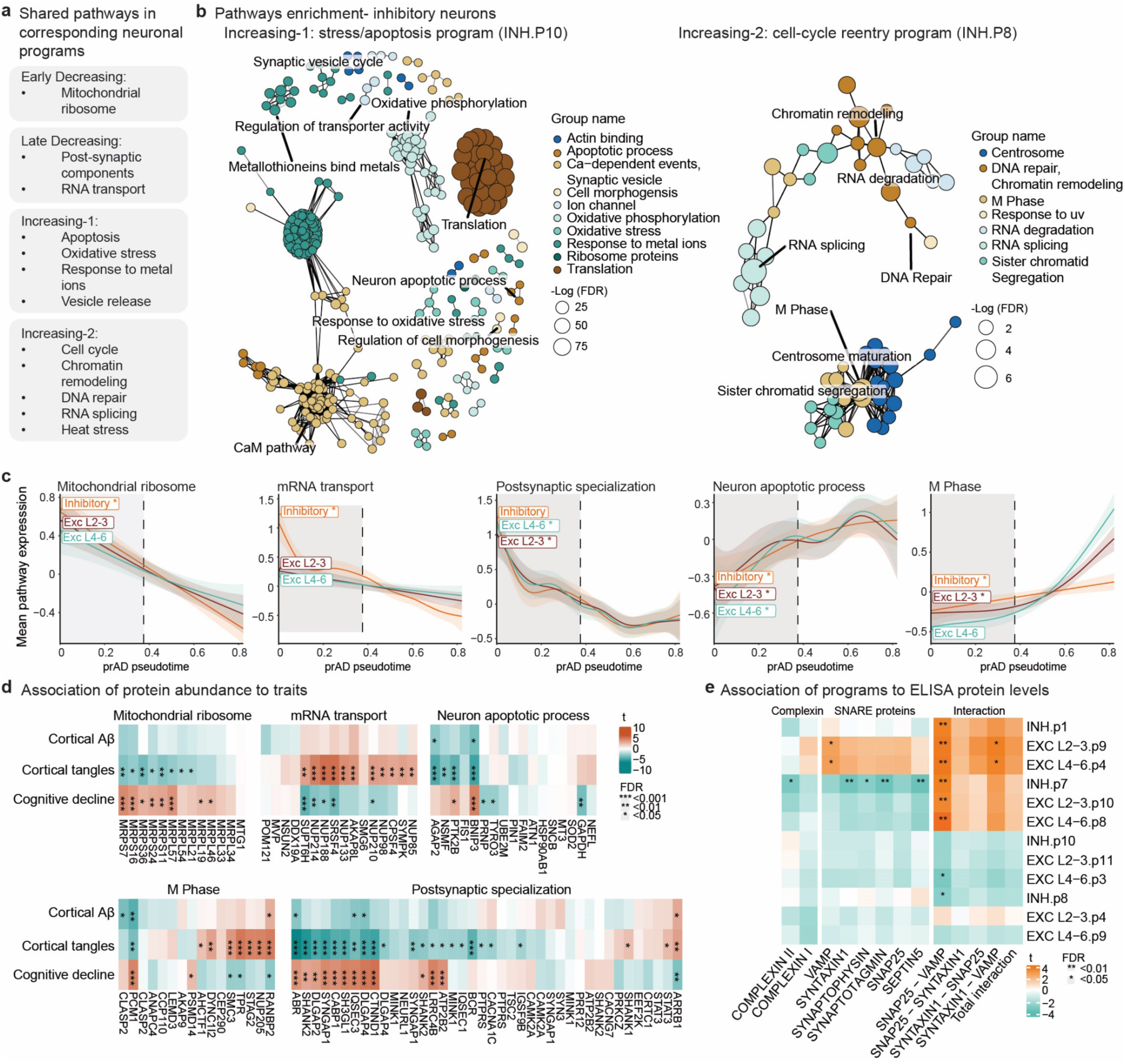
Functional annotation of AD-associated neuronal programs. **a.** Key shared pathways within each group of corresponding neuronal programs between inhibitory, upper layer-excitatory, and lower layer-excitatory neurons. **b.** Overview of clustered enriched pathways revealing distinct biological functions within the inceasing-1 INH.p10 and increasing-2 INH.p8 inhibitory neuronal programs. Nodes = pathways (showing only connected nodes), node size = -log FDR corrected p-value. Edge length and width reflect pairwise pathway distance (correlations over genes assignments to pathways, showing edges only for highly correlated pathways). Colours represent clusters of pathways grouped by overlapping genes and distinguishing pathways with distinct gene compositions. **c.** Dynamics of pathways along the prAD trajectory coordinated across neuronal classes. Inferred dynamics over the pseudotime of averaged, scaled expression of pathway genes identified in the program signatures (**Methods**) across cells per individual. Dashed line represents the split point between trajectories. **d.** Validation of neuronal expression changes in AD by proteomics. Association between proteins and AD traits grouped by biological pathways (n=400, FDR corrected, **Methods**). **e**. Association of ELISA protein levels and interactions of protein pairs to the AD-associated neuronal programs (n=196, FDR corrected, **Methods**).

We validated key differential pathways (**Fig. 3d, Extended Data Fig.5-7**). In an independent snRNA-seq^7^ dataset (**Methods**), we showed down-regulation of the expression of mitochondrial ribosomal and mRNA transport genes, upregulation of DNA repair, M phase genes, and heat stress response genes and a partial upregulation of apoptotic and oxidative stress response genes (**Extended Data Fig. 5b,f, 7a,b**). We further validated the neuronal reprograming at the protein level using bulk proteomics of 400 DLPFC samples from aging individuals from the ROS/MAP cohort^18^, including 146 overlapping donors with the snRNA-seq dataset. We tested the association of protein expression to AD traits in key predicted differential pathways (FDR<0.05), confirming key disease-associated neuronal programs, such as mitochondrial ribosome proteins’ negative association with AD traits and DNA repair proteins’ positive association (**Fig. 3d, Extended Data Fig. 7e)**. For some pathways, the protein expression profile partially aligned with the RNA observations, such as a subset of the oxidative stress response and M-phase proteins (e.g. PRDX1 for oxidative stress and STAG2 for M phase) while others showed no change at the protein level (**Fig. 3d, Extended Data Fig. 6d**). Several pathways were associated with AD traits, confirming these pathways are differentially dysregulated, but we observed an opposite effect on the protein levels compared to the RNA, including nuclear pore complex proteins that positively correlated to tau loads, and heat shock and apoptotic proteins that were negatively correlated (**Fig. 3d, Extended Data Fig. 7e)**.

Moreover, synaptic transmission proteins showed divergent RNA and protein expression: while RNA levels showed decrease in postsynaptic density and specialization genes and an increase in synaptic release genes along the disease progression (**Fig. 3b,c, Extended Data Fig. 5f, Extended Data Fig. 6e-g**), both categories decreased at the protein level, consistent with a recent report^8^ (**Fig. 3d, Extended Data Fig. 6h)**. This RNA-protein discrepancy may reflect a compensatory transcriptional response to synaptic dysfunction and impaired protein translation in AD^19^, though technical effects in resolutions (bulk proteomics vs. single-cell RNA) or changes in neuronal proportions in AD, can’t be excluded.

To further investigate and assess the link between disease-associated neuronal programs and synaptic integrity in AD, we used ELISA measurements of synaptic proteins and complexes in 194 of the snRNA-seq participants, spanning different stages of AD^20^. The ELISA measurements included key synaptic proteins - COMPLEXIN I/II and components of the SNARE complex SNAP25, VAMP, and SYNTAXIN, and their interactions, serving as proxies for functional synapses. We found a strong association between synaptic protein/complex abundance, particularly VAMP and VAMP-SNAP25 interactions, and early-decreasing neuronal programs. In contrast, late-decreasing programs were negatively associated with synaptic proteins levels but positively associated with their interactions, while the increasing programs were negatively associated with the interactions of synaptic proteins (**Fig. 3e**). Coordinated with these findings, synaptic proteins levels showed a significant negative associated with cortical Aβ-load, supporting synaptic impairment early in pre-clinical AD (**Extended Data Fig. 7f**).

### Neurons converge to distinct programs of stress/apoptosis or cell-cycle reentry

The cell-cycle reentry of neurons in AD is highly intriguing and raises questions regarding their role in neurodegeneration. Based on previous reports that observed cell-cycle reentry in aging brains and specifically in AD^21,22^ (with no evidence of cell divisions), we hypothesized that it could either be a step toward apoptosis^23–25^, or could lead to a senescence-like arrest^22,23,26^.

First, we observed that the *cell-cycle reentry programs*, were predominantly enriched for M-phase genes and not for other cell-cycle stages (**Fig. 3**). These programs included markers such as *STAG2* and *CENPC*, involved in sister chromatid cohesion and kinetochore-mediated segregation, and *ANAPC16*, a component of the anaphase-promoting complex. Moreover, we detected early modulation along the AD cascade of DNA damage linked cell-cycle checkpoint genes, including a decrease in *ATR* (activated by DNA damage to regulate cell-cycle progression^27^), *BRCA1* (a cell-cycle checkpoint regulator acting throughout the cell-cycle^28^), *RAD9A* (a cell-cycle checkpoint and DNA damage sensor), along with an increase in *ATM*, which promoted cell-cycle arrest in response to DNA damage^29^. Notably, we found upregulation of *POLK* and *REV1*, key components of the translesion DNA synthesis pathway allow bypass of replication-blocking lesions and ensure DNA replication continuity^30,31^, supporting progression beyond S-phase (Extended Data Fig. 7g).

Next, we sought to dissect the link between the stress/apoptosis and cell-cycle reentry programs. By quantifying their distribution and overlaps across individual cells, we found these two increasing programs to be mutually exclusive. The distribution of signature scores across individual neuronal cells showed these two programs were non-overlapping (**Fig. 4a,b, Extended Data Fig. 8a**) and significantly anti-correlated across cells (**FDR<0.05, Fig 4c, Extended Data Fig. 8b**). Validation in an independent snRNA-seq dataset^7^ confirmed that the apoptosis signature is correlated with oxidative stress, but not with the cell cycle signature across neurons (**Extended Data Fig. 5c)**. Moreover, quantifying the number of neurons expressing each of the two programs (**Methods, Supplementary Table 5**), showed that fewer than 0.3% of neurons co-express both programs **(Fig. 4d, Extended Data Fig. 8d,e)**. Notably, the number neurons expressing *cell-cycle reentry* program increased 3-4 fold in advanced AD compared to healthy individuals, specifically in inhibitory and deep layer (L4-6) excitatory neurons, but not in L2-3 (**Fig. 4e**, **Extended Data Fig. 8f**). This increase was robust across quantification thresholds (**Extended Data Fig. 8g**) and consistent with prior DNA content-based quantification^22^ of 2n/4n neurons in AD and control brains (**Fig. 4f**).

**Figure 4.**
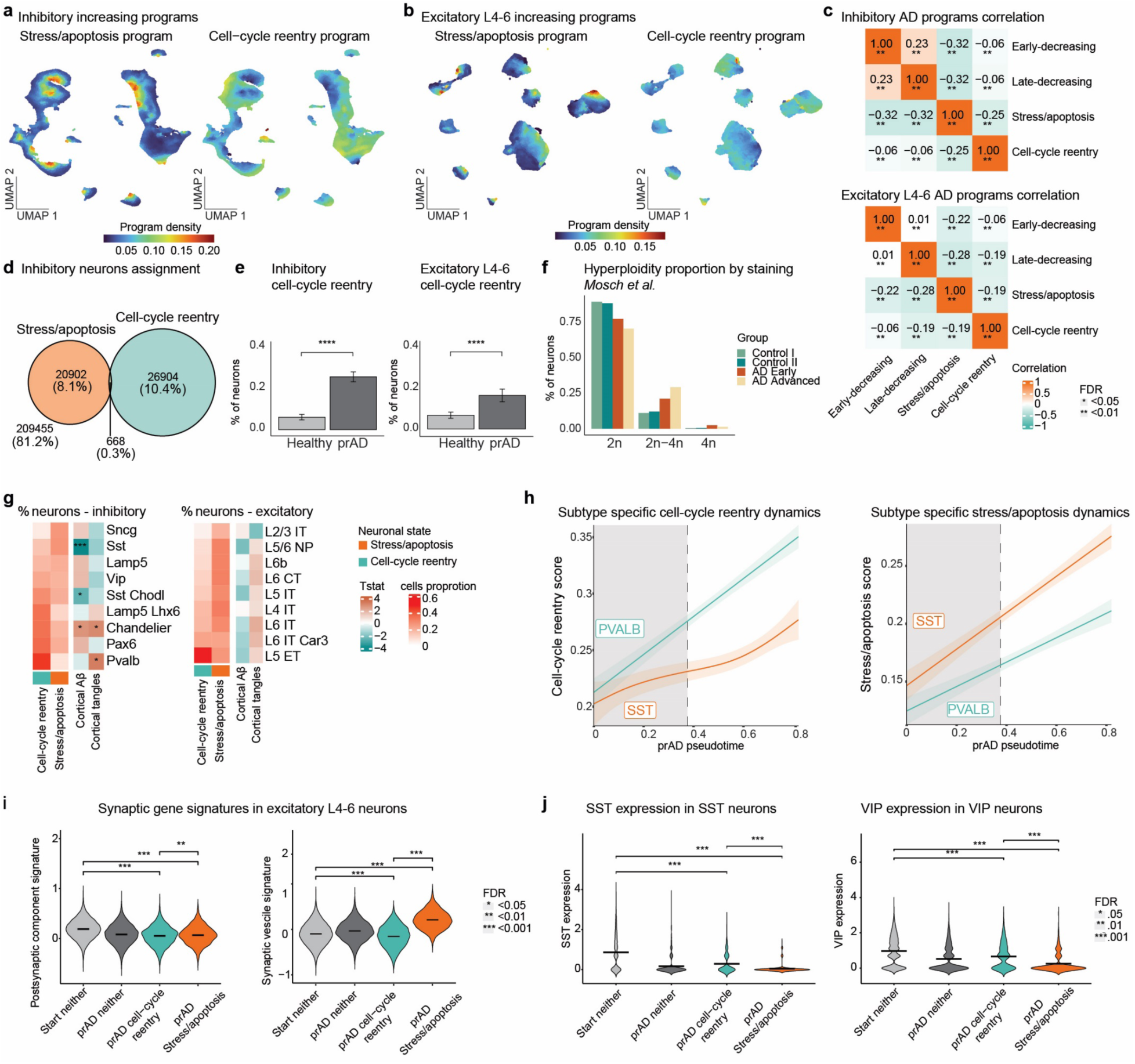
**Mutual exclusive expression of the cell-cycle reentry and the stress/apoptosis neuronal programs linked to neuronal vulnerability. a-d**. Mutual exclusive expression of the cell-cycle reentry and the stress/apoptosis neuronal programs. UMAP embedding of inhibitory (**a**) and excitatory L4-6 (**b**) neurons, coloured by locally smoothed density value of each program per cell. (**c**) Correlations of the decreasing and increasing disease-associated programs across inhibitory (top) and excitatory L4-6 (bottom) neurons. Spearman correlations with FDR-adjusted significance (Excitatory L2-3 in Extended Data Fig. 8a-b). (**d**) Quantification of inhibitory neuronal cells assigned to stress/apoptosis or cell cycle reentry shows minimal overlap (threshold of 95%, **Methods**). **e-f**. Increased cell-cycle reentry neurons in AD. (**e**) Average proportions of cell-cycle reentry neurons in healthy (n=78) and advanced prAD (n=34) groups of individuals (**Methods**). Wilcoxon test p-value<0.001. (**f**) Proportion of hyperploid neurons measured by immunohistochemistry in control and AD brains from Mosch *et al.*^20^. **g-h**. Cell-cycle reentry and stress/apoptosis neuronal programs and linked to resilient and vulnerable neuronal subtypes (respectively). **g**. Inhibitory and excitatory neuronal subtypes ranked by the proportion of cell-cycle reentry and compared to the proportion of stress/apoptosis neurons in advanced AD individuals. Right columns show the association of subtype proportion with cortical Aβ and tau load (FDR<0.05; **Methods**). **h.** Dynamics of cell-cycle reentry (left) and stress/apoptosis (right) average program scores along the prAD trajectory for relative vulnerable SST and resilient PVALB inhibitory neuronal sub-types. **i**. Postsynaptic genes and presynaptic vesicles averaged scaled expression in excitatory L4-6 cell groups, cells from healthy individuals (n=46272), cell-cycle reentry (n=9601), stress/apoptosis (n=2627) or neither (n=3735) in advanced AD. Wilcoxon test FDR<0.001. (j) Neurotransmitters SST and VIP expression within inhibitory neurons of healthy individuals (SST n=10832, VIP n=10524), cell-cycle reentry neurons (SST n=914, VIP n=485), stress/apoptosis (SST n=598, VIP n=667) or neither (SST n=10832, VIP n=10524) in advanced AD individuals. Wilcoxon test FDR<0.001.

Overall, our analysis suggests the two programs represent distinct neuronal fates in AD, rather than sequential phases of a single process, supporting that cell-cycle reentry in neurons may lead to arrest at M-phase and not to apoptosis. However, we cannot exclude the possibility that some cell-cycle reentry events may progress to cell death at or before M-phase, but are too transient to be robustly captured.

### The increasing programs of stress/apoptosis or cell-cycle reentry align with neuronal vulnerability

Comparison of program abundances across neuronal subtypes, each linked to a different degree of vulnerability by its negative association to the Aβ and tau load (**Methods**), revealed a general pattern: Vulnerable neuronal subtypes, whose proportions decrease faster as disease progresses, such as SST, SNCG and LAMP5 interneurons and L2/3 IT pyramidal neurons, had higher abundance of cells expressing the stress/apoptosis program; In contrast, resilient neuronal subtypes, such as PV interneurons and L5 ET, had higher abundance of cells expressing the cell-cycle reentry program (**Fig. 4g**). Furthermore, comparing the dynamics of the programs between subtypes (fitting subtype-specific patterns, **Methods**), showed that while both programs start at the same level and increase in all neuronal subtypes, the vulnerable SST interneurons consistently express higher levels of the stress/apoptosis program whereas the resilient PV interneurons consistently express higher levels of the cell-cycle reentry program along AD progression (**Fig. 4h)**. In contrast, the dynamics of decreasing programs did not differ between neuronal subtypes (**Extended Data Fig. 8h**).

Next, we aimed to gain deeper insight into the functional differences between stress/apoptosis and cell-cycle reentry neurons. As an indication of impaired or preserved synaptic functions, we compared the expression levels of neurotransmitters, pre-synaptic vesicle release and structural post-synaptic genes between stress/apoptosis vs. cell-cycle reentry neurons in advanced AD brains, and neurons from healthy individuals (**Fig. 4i,j**). Stress/apoptosis neurons exhibited the most pronounced change in presynaptic genes, whereas cell-cycle reentry neurons maintained expression patterns closer to healthy cells (**Fig. 4i**), including the expression of the neurotransmitter genes SST and VIP within the respective subtypes (**Fig. 4j**). Moreover, neurotransmitter expression negatively correlated with the stress/apoptotic program score but not with the cell-cycle reentry (**Extended Data Fig 8i)**. Reduced expression of post-synaptic genes was observed for both neuronal groups **(Fig 4i)**. The findings suggest that the cell-cycle reentry neurons may maintain synaptic functions at least in part, in contrast to stress/apoptosis neurons.

Together, these results reinforce the notion that AD neurons adopt one of two broad fates - a pro-apoptotic stress program vs. a possible survival-promoting cell-cycle reentry program. The balance between these two programs may underlie the observed neuronal subtype-specific vulnerability levels.

### Coordinated neuro-glia programs along the AD cascade

We next investigated the extent of neuronal and glial coordination, by mapping program dynamics along the prAD and ABA trajectories and their co-occurrence by the BEYOND framework^6^ (**Extended Data Fig. 9a, Methods**). We revealed highly coordinated neuronal and glial programs (**Fig. 5a,b, Extended Data Fig. 9b,c**), partitioned into three communities of coordinated neuro-glial programs with shared prevalence across individuals and similar dynamic patterns along the disease and aging trajectories (**Fig. 5a, Extended Data Fig. 8c, Methods**).

**Figure 5.**
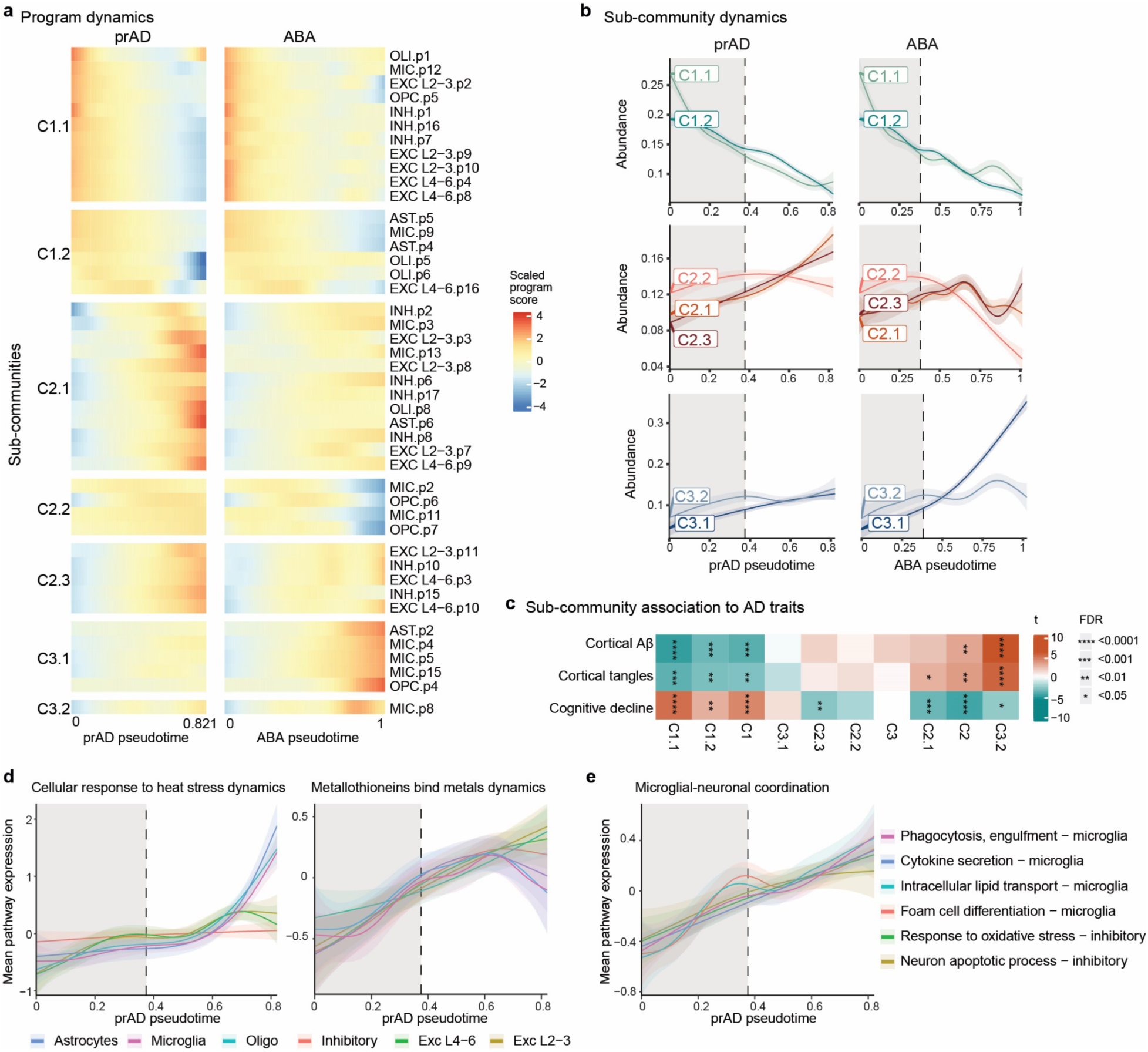
Coordinated glial neuronal programs along AD progression. a-b. Communities of coordinated programs in glial and neurons in aging brains. **a.** Programs dynamics along the prAD pseudotime (left) and the ABA pseudotime (right) divided into 7 sub-communities (clustered by BEYOND^6^, **Methods**). Colour by the inferred program abundance scaled along the pseudotime. **b**. Distinct dynamic patterns of sub-communities along the two trajectories (prAD, left; ABA, right). Inferred dynamics over averaged of programs scores within the community (**Methods**). Dashed line represents the split point between trajectories. **c.** Associations between proportions of sub-communities and AD traits (linear regression *t*-test, FDR < 0.05, **Methods**). **d**. Similar dynamics of the same pathway across different cell types along the prAD pseudotime. Inferred dynamics of averaged pathway scores per individual within the cell type along the prAD trajectory (**Methods**). Left: Cellular response to heat stress, right: Metallothioneins bind metals. **e**. Distinct pathways with similar dynamics across neurons and microglia cells along the prAD pseudotime (as in d).

Community C1 contained the *homeostatic* programs, decreasing in AD and aging, including homeostatic glial programs and the decreasing neuronal programs (**Fig. 5a),** indicating early parallel changes in glial and neurons. Community C2 contained the AD-associated programs, increasing predominantly along the prAD trajectory, which were further sub-clustered into three sub-communities: the disease/stressed-glial (OLI.p8, AST.p6, and MIC.p13) and cell-cycle reentry neuronal programs - community C2.1; the ABA-only decreasing programs – community C2.2; and the stress/apoptosis neuronal community - C2.3. Community C3 contained programs increasing predominantly along the ABA trajectory, including reactive glial programs matching previous reports^6^ (**Fig. 1e** and **Extended Data Fig. 2c,i**). Interestingly, our analysis distinguishes between the DAM-like immune-activated MIC.p8 that increase in both prAD and ABA, versus the foam-lipid associated MIC.p11, which is unique to the prAD trajectory (**Fig. 5a**). Associating the abundance of the cellular communities to AD-traits (**Methods**) revealed that the homeostatic community C1 was negatively associated with disease pathologies, Aβ and tau, while the disease community C2 was positively associated with disease pathologies and negatively with the cognitive slope (FDR<0.05, **Fig. 5c**).

Pathway analysis revealed coordinated neuro-glial responses along AD progression. Stress responses, specifically response to heat stress and to metal ions increased across neurons and glia with AD progression (**Fig. 5d**), with response to heat stress appearing earlier in neurons compared to glia, suggesting greater early vulnerability (**Fig. 5d**). Alignment of cell-type specific functions, aligned neuronal increase in oxidative stress and apoptotic genes with microglial activation of foam-cell differentiation, lipid transport, cytokine secretion, and phagocytosis (**Fig. 5f**). Overall, these findings show early modulations of neurons in AD that are tightly coordinated with altered glial functions, with both shared and cell-type specific responses that might react to and further affect neuronal impairments.

### Neuronal and glial programs distinguish AD from alternative aging

We next show that the corresponding AD-associated neuronal programs distinguish between AD and alternative brain aging (ABA) along with unique glial responses in AD compared to ABA. Comparing the dynamics of neuronal programs along the two trajectories of prAD and ABA (**Fig. 2d**), we found in the ABA only a moderate or no decrease of the early and late decreasing programs, a significantly lower increase of the stress/apoptosis programs, and a moderate increase in the cell-cycle reentry programs (**Fig 6a**). Both the late-decreasing and the stress/apoptosis increasing programs showed high variability in the ABA trajectory, indicating potentially additional unresolved variability in this aging trajectory (**Fig 6a**). Notably, the program dynamics along published trajectories of AD and aging^6^, validated the specificity of the decreasing and increasing neuronal programs to the prAD trajectory showing no change in the ABA trajectory (**Extended Data Fig. 9d**).

**Figure 6.**
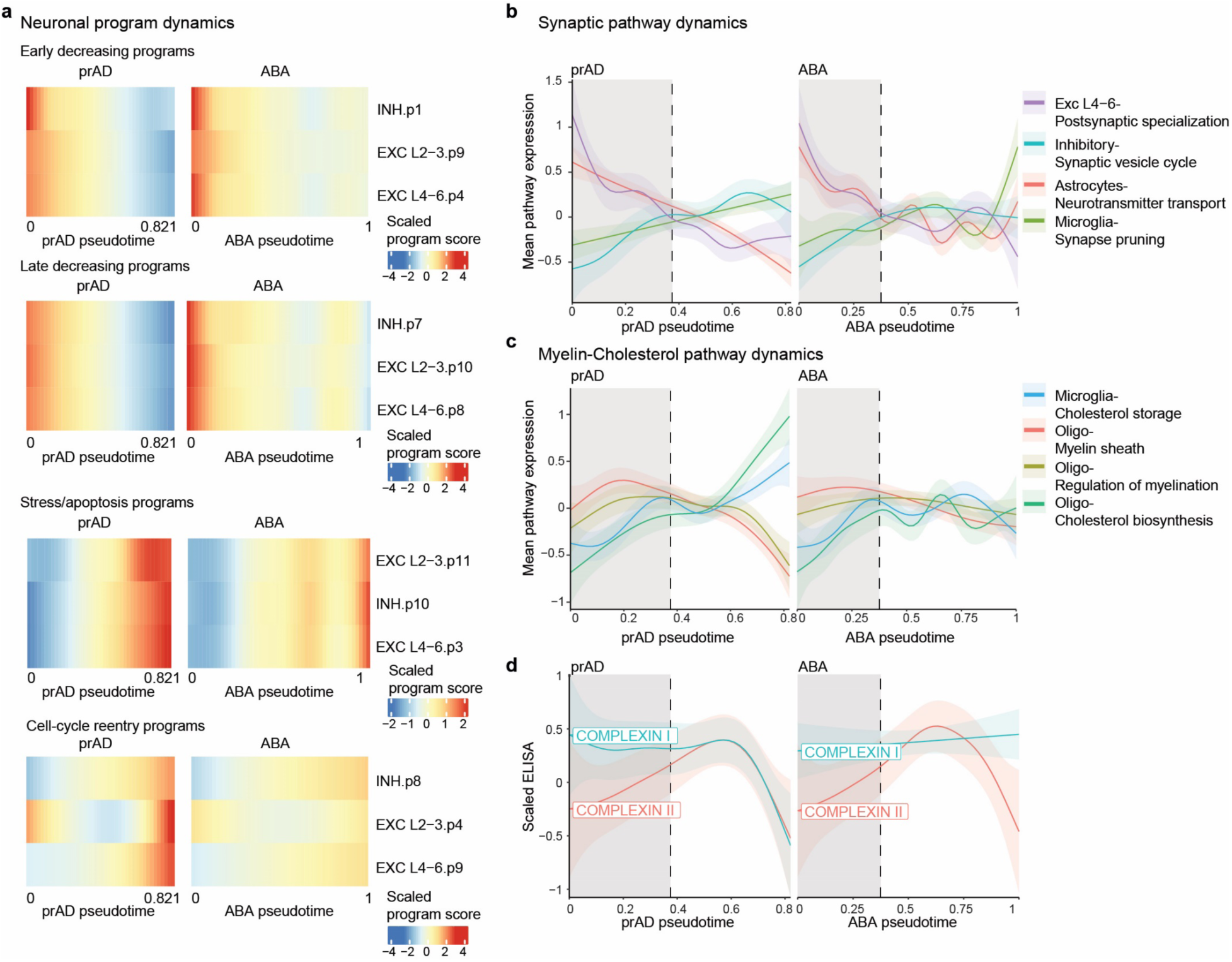
Distinct neuronal and glial programs in prAD and ABA. **a.** AD-associated neuronal programs are strongly modulated in the prAD trajectory compared to the ABA. Dynamics of the groups of corresponding decreasing and increasing neuronal programs along the prAD trajectory (left) and the ABA trajectory (right). Colour represents the level of averaged program score inferred per pseudotime, scaled across both trajectories. **b.** Coordinated synaptic changes are unique to the prAD trajectory. Inferred dynamics of averaged pathway scores per individual within the cell type along the prAD and ABA trajectories. **c.** Myelin-cholesterol expression dynamics in oligodendrocytes and microglia are unique to the prAD trajectory. Presented as in b. **d.** Reduction of COMPLEXIN I is unique to the prAD trajectory. Dynamics of COMPLEXIN I and COMPLEXIN II proteins levels measured by ELISA (n=192) along the prAD and ABA pseudotime. Dashed line represents the split point between trajectories.

At the pathway level, we found coordinated modulation of synaptic-associated pathways in neurons and glia as well as changes in myelin production pathways to be unique to the prAD, without any change along ABA progression (**Fig. 6b,c**). Aligning expression dynamics of pathways across neuronal and glial cells in the prAD and ABA trajectories, uncovered in the prAD a coordinated decreased expression of postsynaptic genes by neurons and neurotransmitter transport genes by astrocytes, and a coordinated increased expression of presynaptic genes in neurons accompanied by synaptic pruning genes in microglia, with no clear change in the ABA trajectory (**Fig. 6b**). Furthermore, we found a sharp decreased expression of myelin production and regulation genes in oligodendrocytes and a sharp increase of cholesterol-related pathways by oligodendrocytes and microglia, aligning with reports of impaired myelin and cholesterol pathways in AD^7,32,33^, with no such changes recorded in the ABA trajectory (**Fig. 6c)**.

The relatively late changes in myelin and cholesterol pathways, consistent with previous reports^6^, appear to follow earlier coordinated neuronal and glial changes, including synaptic gene modulation, yet their sharp transition marks a dramatic shift in brain state at this disease stage. To align this transcriptional timeline with synaptic impairment, we examined the dynamics of synaptic protein expression using ELISA measurements^20^. Most strikingly, COMPLEXIN-I, a key regulator of neurotransmitter release, showed a sharp decrease specifically along the prAD trajectory (**Fig. 6d**), indicating more pronounced synaptic damage compared to ABA. This decline aligned with the sharp changes in cholesterol and myelin-related genes (**Fig. 6c,d**), as well as the rise in glial heat stress responses (**Fig. 5d**). Together, these converging dynamics mark a critical phase in AD progression, characterized by coordinated neuronal and glial stress, synaptic loss, and myelin dysregulation.

## Discussion

Here, we provided a comprehensive view of the early molecular events that drive AD progression, revealing a coordinated, transcriptional reprogramming of neurons – shared across all subtypes and classes - that emerges early along the disease cascade, well before clinical cognitive decline. These neuronal changes are tightly coupled to glial responses, underscoring a neuron–glia interplay that is specific to the AD cascade and distinct from alternative brain-aging trajectories.

Importantly, our study adopts a continuous-modeling framework for sc/snRNA-seq analysis. By representing each cell as a weighted combination of transcriptional programs, we captured nuanced and partially overlapping cellular functions that clustering methods often miss. While an alternative exhaustive sub-clustering approach might also partition cells by functions, it is unlikely to fully resolve intersecting axes of variation; by contrast, the continuous approach required far fewer cells and is better suited to capture the multifaceted nature of cellular identity and functions. This approach uncovered neuronal subtype–specific as well as shared responses across subtypes including coordinated disease responses, and readily detected subtle state transitions in glia. For example, we separated the DAM-like immune program^16^—elevated in aging and AD—from AD-specific foam-lipid programs^6^ in microglia, providing mechanistic insights and clearer biomarkers for follow-up studies. This partition is consistent with efforts to define microglial factors in an independent dataset^34^. The programs identified previously reported disease and aging subpopulations^4–8^, and the inferred trajectories and pseudotime agree with published cluster-based results^6^, reinforcing this framework’s robustness to model changes in cellular environments.

A key finding is the existence of two mutually exclusive neuronal programs in AD. One, enriched in vulnerable subtypes, combines expression profiles of cellular-stress and apoptotic signatures and coincides with the dysregulation of synaptic gene expression, suggesting synaptic dysfunction and potentially explaining the selective neuronal loss in these subtypes. The other, combines expression of DNA-damage responses, heat-shock signaling, and cell-cycle M-phase genes, and predominates in more resilient neuronal subtypes. We cannot exclude the possibility that a subset of cells reenters the cell-cycle and subsequently undergoes apoptosis, as previously suggested^23–25^. However, as our data as well as prior immunohistochemistry-based estimates^22^ suggest that approximately 20% of neurons in advanced AD are in M phase, and the abundance of this program in resilient neuronal subtypes, indicate that cell-cycle reentry may lead to arrest and promote survival, rather than trigger apoptosis. Moreover, though functional analyses of post-mortem tissue are not possible, the transcriptional signatures indicate that cell-cycle arrested neurons may retain synaptic functions, as they express synaptic genes at a level closer to those of healthy neurons compared to stress/apoptotic neurons.

Previous studies have shown that neurons, although post-mitotic, can reenter the cell-cycle with age, a process exacerbated in AD^21,22^ and potentially induced by DNA damage^23^. However, the regulatory mechanisms governing this transition remain unclear, particularly the ability of neurons to undergo DNA replication despite damage, which would typically trigger apoptosis. Here, we show that early neuronal reprogramming includes extensive changes in epigenetic regulators and specific dysregulation of DNA damage checkpoints that could underlie the cell-cycle reentry and escape from apoptosis. Such changes include downregulation of BRCA1, a key sensor of DNA damage that promotes apoptosis^28^, along with the induced expression of key components of the translesion DNA synthesis pathway (POLK and REV1) that enable bypass of DNA replication-blocking^30,31^.

Mapping neuronal and glial programs along disease progression clarified the relative timing of their responses along disease progression and the deviation from the aging process. An early, global decline in glial homeostatic programs, as previously observed^5,6^, is accompanied by parallel fast decrease in neuronal mitochondrial–ribosomal genes, and slower subsequent synaptic-gene modulation in neurons. We aligned specific function across cell types along the disease timeline, such as the suppression of neuronal synaptic transmission genes expression with the reduced neurotransmitter-transport pathways in astrocytes that together underlie the observed synaptic dysfunction in neurons in AD^35^. In particular, consistent with stress-responding Ast.10 predicted as a driver of AD in our earlier work^6,36^, stress-response pathways rise across cell types, though neurons mount an initial stress wave, followed by a shared response by all glia cells, supporting the increased vulnerability of neurons as early drivers of the AD cascade^37–39^. We uncovered AD-specific cellular modulations, highlighting synaptic associated changes in neurons, foam-cell lipid metabolism in microglia (MIC.p11), stress responses in astrocytes (AST.p6), and myelin impairment in oligodendrocytes (OLI.p5-6) as key pathways differentiating the AD cascade from alternative brain aging. Moreover, we found that the changes in myelin and cholesterol occur at later disease stages, yet they mark the entry of a critical phase of synchronized neuronal and glial deterioration and accelerated synaptic loss.

Overall, we show that from the earliest stages of AD, neurons undergo extensive reprogramming in concert with loss of critical homeostatic-supportive functions in glia cells and increased neurotoxic pathways. Consequently, neurons in AD either enter apoptosis following severe stress, oxidative and others, or alternatively, reenter the cell-cycle and arrest at M-phase, a program associated with improved survival. These insights redefine neuronal vulnerability, illuminate mechanisms of resilience, and highlight new therapeutic entry points for intervention before overt cognitive decline.

## Methods

### Dataset description: Study participants, AD traits, and snRNA-seq dataset

The cell-type specific gene programs atlas was derived from an snRNA-seq dataset^6^ of 1.58 million single nucleus RNA profiles of neuronal and glial cells from 437 Dorsolateral Prefrontal Cortex (DLPFC) samples from participants of the aging ROS/MAP cohorts. The snRNA-seq dataset was fully annotated to cell types and cell sub-populations by clustering analysis in our previous work published in Green et al^6^. Additional datasets used for validations and annotations, include bulk proteomics of 400 individuals^18^, with 146 overlapping the snRNA-seq dataset, and by ELISA measurements of synaptic proteins in 194 individuals^20^ overlapping the snRNA-seq dataset.

The snRNA-seq profiled participants were selected blind to their neuropathologic and clinical traits, and based on availability of frozen pathologic material from the DLPFC (BA9) brain region, including only participants with RIN>5 and post mortem interval (PMI) <24 hours. The study cohort includes diverse individuals across the full range of the pathological and clinical stages of AD. The demographic and clinicopathologic characteristics are described in **Supplementary Table 1** (and in Green et al^6^).

Each individual profiled in these datasets was enrolled in one of two longitudinal clinical-pathologic cohort studies of aging and dementia, the Religious Orders Study (ROS)^14^ and the Rush Memory and Aging Project (MAP)^15^, collectively referred to as ROSMAP. All participants are without known dementia at enrolment, have annual clinical evaluations, and agree in advance to brain donation at death. At death, the brains undergo a quantitative neuropathologic assessment, and the participant’s rate of cognitive decline is calculated from the longitudinal cognitive measures that include up to 25 yearly evaluations^41^. Each study was approved by an Institutional Review Board of Rush University Medical Center. All participants signed an informed consent, Anatomic Gift Act, and repository consent.

Pathological measures were collected as part of the ROS/MAP cohorts (previously described in details^42–44^). We focused our analysis on three quantitative AD-related traits, which have a greater statistical power compared to discrete classifications: **Rate of cognitive decline**: Uniform structured clinical evaluations, including a comprehensive cognitive assessment, are administered annually to the ROS and MAP participants. The ROS and MAP methods of assessing cognition have been extensively summarized in previous publications^45–47^. Scores from 19 cognitive performance tests common in both studies, 17 of which were used to obtain a summary measure for global cognition as well as measures for five cognitive domains of episodic memory, visuospatial ability, perceptual speed, semantic memory, and working memory. The summary measure for global cognition is calculated by averaging the standardized scores of the 17 tests, and the summary measure for each domain is calculated similarly by averaging the standardized scores of the tests specific to that domain. To obtain a measurement of cognitive decline, the annual global cognitive scores are modelled longitudinally with a mixed effects model, adjusting for age, sex and education, providing person specific random slopes of decline (which we refer to as ***cognitive decline***). Further details of the statistical methodology have been previously described^48^. **Aβ load and tau** (cortical paired helical filaments tau) **tangle density**: Quantification and estimation of the amount of parenchymal deposition of Aβ and the density of abnormally phosphorylated tau-positive neurofibrillary tangles levels present at death (which we refer to as Aβ and tau pathology, respectively). Tissue was dissected from eight regions of the brain: the hippocampus, entorhinal cortex, anterior cingulate cortex, midfrontal cortex, superior frontal cortex, inferior temporal cortex, angular gyrus, and calcarine cortex. 20µm sections from each region were stained with antibodies for the Aβ and tau protein, and quantified with image analysis and stereology. Measurements were summarized to provide a global measure of Aβ and tau loads. For the trait association, causality prediction and dynamics modelling analyses, we used the measurements of Aβ and tau in the midfrontal cortex (referred to as cortical Aβ and -tau). The load of these pathologies at the midfrontal cortex are a potentially better proxy for the pathology in the DLPFC brain region compared to the pathology load across the entire brain. Further, we note that we are assessing cellular changes in the fresh-frozen DLPFC samples from one hemisphere of each brain in relation to the measures of cortical Aβ and tau loads measured in the midfrontal cortex of the opposite, fixed hemisphere in which the standard, structured neuropathologic assessment is conducted.

### Constructing an atlas of Cell-type specific gene expression programs by Topic modelling

#### Construction of the gene program atlas

To model the continuous diversity in aging human brain cells, we used the snRNA-seq dataset of 1.58 million cells captured from 437 aging individuals, and the cell type classification done in Green at el^6^ we separated the cells to the different types and fitted co-expression gene programs separately for each cell type. Co-expression gene programs were modelled using the fast-topic algorithm^13^ based on Non-negative Matrix Factorization (NMF) to identify co-expressed programs of genes (called topics), and model cells as a linear combination of programs (topics). Each cell is assigned a score for each program, and each program assigns a weight for each gene.

The gene expression matrix *x_n_*_x*m*_ is factorized into two non-negative matrices – *F_n_*_x*k*_ and *L_m_*_x*k*_, with no negative elements, such that approximately *X* ≈ *FL^T^*, where: n=number of genes; m=number of cells; k=number of programs; *X_ij_* = The expression of gene i in sample j; *F_ij_* = the weight of gene i in program j; *L_ij_* = the score of program j in cell i.

For each cell type, we fitted a range of a number of programs (k) using the fastTopics^13^ package (version 0.6-135, fit_topic_model). The final number of topics was decided by the alignment of programs with the literature showing the number of programs is sufficient to capture (at least) known subtypes and states and yet is not too large such that each program is found across multiple individuals and has a unique signature of differential genes (discussed more in the next section). Validations of programs robustness were done by fitting programs over a random subset of cells or different number of programs, and comparing the programs scores across cells. We compared the fit of programs with a different number of programs or across subsets of cells, by computing the Spearman correlation between the program scores (cor, method = “spearman”). P-values were corrected for multiple hypothesis testing using Benjamini–Hochberg procedure. We validated the robustness of our conclusion, showing consistent results in downstream analysis of trait associations, trajectories and dynamics for different number of programs, to ensure that robustness of the conclusions was not derived from the chosen number of topics.

The neuronal cells were split to three classes and we fitted the program separately for each: inhibitory neurons, upper cortical layer L2-3 neurons (*CUX2*^+^ nuclei), and deep cortical layers L4-6 (*CUX2*^+^ nuclei). Glial cell types were split to: microglia, astrocytes, oligodendrocytes and oligodendrocyte precursor cells (OPCs). We excluded the vascular niche cells because of the lower number of cells per sample compared to the large cellular diversity within them.

For validations of the shared neuronal programs, inhibitory neurons were further split to subtypes by clustering annotated by neurotransmitters and markers expression, and the same methods were applied to each subtype separately. The subtype-derived programs were compared by the overlap of gene signatures using the Jaccard similarity index to test the similarity of programs inferred across subtypes.

#### Differential expression and functional annotations

For each of the programs in each cell type, we assigned a signature, based on the set of differentially expressed genes using fastTopics^49^ (de_analysis, lfc.stat = vsnull) and filtered the upregulated genes (postmean > 0) with lfsr < 0.05. The list was then filtered to include only the program-specific genes. For this end, we establish a weighting scheme that takes as input the program-gene scoring matrix F(n x k) matrix (F_ij_ = the weight of gene i in program j, as described above) and produces a new, reweighted F’ matrix. This scheme prioritizes genes that are uniquely highly expressed in each topic, while enabling overlapping genes across programs. The process is as follows: For the *i*-th gene and the *j*-th program we define an upper (0.95 and 0.8 quantiles for neuronal cells and glial cells, respectively) and lower quantiles threshold (0.5 quantile). First, we filtered all of the genes that were lower than the upper quantile then we computed the reweighted score matrix:

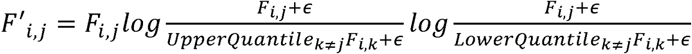

Consequently, all genes within a program that had a higher score compared to the upper quantile threshold across all programs, will receive a positive and high reweighted score. Specifically, we seek the new scores to be higher, as the log ratio of their scores within the program compared to other programs is higher (indicative of uniqueness), and taking into account the original scores indicative of the relevance of their expression levels to the program. While genes that are not unique to the program, their reweighted score will be reduced compared to the original score. A z-score is computed over the log of the reweighted score. To filter down and increase our confidence in the differential expressed genes of each program, we filter the fastTopics output by taking only the genes with z-score >1.

Functional annotations were inferred by (1) identifying statistically significant enriched pathways and gene sets in the differential gene signatures and (2) grouping enriched pathways based on their gene-similarities to reflect distinct functions rather than a list of overlapping annotated gene-sets. We identified enriched pathways by hyper-geometric test using the clusterProfiler^50^ tool (compareCluster, formula=id∼cluster+direction) over the gene-set databases: KEGG (fun=enrichKEGG, organism=hsa), Reactome (fun=enrichPathway) and GO (fun=enrichGo, OrgDb=org.Hs.eg.db, ont=BP, MF, CC). Correcting for multiple hypothesis by FDR with a threshold of p.adjust(Benjamini Hochberg correction)<0.05 and q-value<0.2, setting the background gene set as the all genes measured in the snRNA-seq data as expressed within the relevant cell type. Redundancy between GO terms were removed by clustering using ClusterProfiler (simplify function). To better understand the functionality captured by the pathways and overcome the issue of redundancy within the databases—where multiple different enriched pathways capture the same sets of differential genes and therefore probably reflect the same function—we clustered the enriched pathways of each program based on their overlapping genes. Given a list of enriched pathways, we first constructed a binary matrix of genes assignment to pathways – for each pathway, all genes included in the pathway that are also differentially expressed in the program are marked as 1, and those not included as 0. Then we applied hierarchal clustering of pathways based on their degree of overlap of genes (method hclust, clustering_distance=correlation). For visualization we plotted the enriched pathways and their overlap as a graph, where each node is a pathway, and edges connect between pathway with high gene overlaps, specifically using correlation threshold of 0.3 for the enriched pathways of program INH.p8 and correlation threshold of 0.7 for program INH.p10 (due to the large number of enriched pathways in that program).

#### Antisense gene enrichment

To test for enrichment of gene programs in antisense genes, we identified antisense genes using the following regular expression “.*-AS\d+$”. We then used hyper geometric test (stats package, *phyper*) with all genes measured in the cell type as the background. We carried the analysis for all programs inside each cell type and corrected for multiple-hypothesis testing by calculating the FDR (<0.05, p.adjust, method=BH) within each cell type.

#### Programs similarity

To test similarities of the neuronal programs across the inhibitory, upper layer excitatory (L2-3) and lower layer excitatory (L4-6) neuronal classes we used two approaches. One, to measure the coordination of the programs across participants we computed the pairwise spearman correlation matrix of the abundance of neuronal programs, defined as the mean program score across all cells per participant, see Matrix P defined below (*cor* method, stats package, use=pairwise.complete.obs, method=spearman). The results were corrected for multiple-hypothesis testing by calculating the FDR (<0.05, p.adjust, method=BH). And second, to compare the gene signature of each program by computing the Jaccard similarity index between the gene signature of each program.

### Association of cell-type specific gene expression programs to AD traits

#### Computation of expression programs abundance

For each individual, we computed its <u>proportions (referred to as abundance) by its</u> mean program score within each cell type or neuronal class, for each of the 87 programs within our atlas. Since the sum of all program scores per cell is one, the mean per individual of all programs found for each cell type or neuronal class also sum up to 1. Then, we unified all the expression program abundance to create the program abundance matrix whose rows represent the participants and columns represent expression programs:

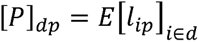

For, *d* = participant, *p* = program and *l* = the loading matrix of the expression program in a certain cell type. We refer to the row *p_d_* as the cellular environment of participant *d*. A column *p_p_* is the vector of mean program score for a program across all participants. Matrix P was used for association statistical analysis as well as the dynamic inference by BEYOND analysis (see sections below).

#### Trait associations statistical analysis

Statistical associations between traits and programs were tested by regressing traits on the square-root abundance of the program:

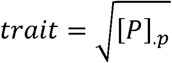

for program *p* and P the abundance matrix (defined above). To remove potential confounding effects, we adjusted for age at death, sex and post-mortem interval (PMI). The results were corrected for multiple-hypothesis testing by calculating the FDR values (<0.05, p.adjust, method=BH) within each tested trait. The trait association analysis was carried out to the AD-traits (described above): cortical Aβ, cortical tangles, cognitive slope, as well as to synaptic proteins levels measured by ELISA.

We also carried out similar statistical associations between AD traits to protein levels measured by proteomics and specific synaptic proteins measured by ELISA for validation. To measure the degree of neuronal subtype vulnerability, we tested the statistical associations between AD traits to the proportion of each neuronal subtype across individuals.

### Validations by independent snRNA-seq dataset, proteomics, and ELISA

#### Proteomics analysis

To validate transcriptomic measurements at the protein level we used 400 proteomics samples from the DLPFC (BA9) of aging post-mortem brains from healthy and Alzheimer’s disease patients^18^ (**Supplementary Table 1)**. Data measurements by label-free quantitation mass spectrometry, capturing 8817 proteins across all samples, and the protein expression levels are scaled across samples as previously described in ^18^. Among the 400 profiled individuals, 146 donors are shared with the snRNA-seq dataset. As proteomics data measured bulk tissue levels of proteins, whereas snRNA-seq measured expression with single-cell resolution, it is a challenge for direct comparisons. To address this issue, we computed the correlation between the expression of each gene of interest at the single-cell level (averaged across all cells of the relevant cell type) and its corresponding expression in the bulk RNA-seq profile from the same donor. This approach enables us to assess whether observed changes in protein levels are more likely to be associated with the cell type of interest or influenced by other cell types present in the tissue. Next, we computed the association between the protein levels and AD traits (as described above).

#### SEA-AD snRNA-seq dataset analysis

In order to validate our results in an independent snRNA-seq dataset, we used the SEA-AD dataset (Gabitto *et al*^7^) taken from an independent aging brain cohort. We used the differentially expressed genes computed over the continuous pseudoprogression score over disease pathologies (effect_sizes.h5ad and pvalues.h5ad from https://sea-ad-single-cell-profiling.s3.amazonaws.com/index.html#MTG/RNAseq/Supplementary <u>Information/Nebula Results</u>). Filtering the subtypes and genes with corrected p-value below 0.05. In each neuronal class (inhibitory and excitatory), we counted the number of subtypes that are positively associated for each gene, denoted *#positive* and the number of subtypes that are negatively associated with the gene, denoted *#negative*. We calculated the difference : *#positive* - *#negative*, Therefore, a positive number indicates that more subtypes upregulated this gene, and a negative number indicates that more subtypes downregulated the gene.

For co-expression validation of biological pathways we computed the signature score of all the differentially expressed genes of each pathway (scanpy, score_genes) and computed their Spearman correlation over the cells (cor.test, method=spearman) and corrected to FDR<0.05 p.adjust, methods=BH).

#### ELISA analysis

To investigate and assess the link of the disease-associated neuronal programs to synapse integrity in AD, we used ELISA measurements of synaptic proteins and complexes in 194 donors from the MAP cohort shared with our snRNA-seq dataset from the MAP cohort along different stages of AD^20^. The dataset included key synaptic proteins, COMPLEXIN I/II as well as the components of the SNARE complex (SNAP25, VAMP, and SYNTAXIN), and their interactions, as an indication for functional synapses. We applied trait associations statistical analysis (as described above) between the abundance of each neuronal program and the proteins levels measured by ELISA. Additional, analysis was done between ELISA protein levels and AD traits.

We also fitted the dynamics of the protein’s levels along the two aging trajectories and pseudotime by GAM (as detailed bellow).

### Assigning samples to trajectories of aging and inference gene expression dynamics and communities of coordinated programs

#### BEYOND analysis – Inference of aging trajectories and pseudotime

To model the dynamics of programs and genes along the disease development and disentangle the disease progression from other forms of aging we adapted the recent BEYOND framework (as described in *Green et al*^6^) that models trajectories of cellular change and assign samples to trajectories and specific pseudotime along the trajectory from information of expression programs (instead of the relative prevalence of discrete cell clusters). We used the mean gene expression programs per participant over all cell types (total of 87 programs). The BEYOND strategy is composed of four major steps: (1) learning the cellular landscape manifold; (2) identifying axes of cellular change (*i.e. trajectories*); (3) fitting programs and trait dynamics along each trajectory; and (4) grouping gene expression programs into coordinated responses across cell-types called *communities*. To implement this strategy, we used the mean program expression matrix aggregated across cell types (see abundance matrix P, defined above). We represent a participant’s cellular environment by its compositional profile of gene expression programs from all analyzed cell types (that is, matrix rows). The cellular landscape is thus represented by the vector space spanned by the columns of the matrix. For convenience, we stored the abundance matrix as an AnnData^51^ object.

Specifically, we applied the following steps:

<u>Step 1: learning the cellular landscape manifold:</u>

We embedded individuals represented by their cellular environments – the diversity of expression programs in neurons and glial - using two manifold learning algorithms for visualization: (1) 2D PHATE embedding (scanpy, external.tl.phate, k=15, n_components=2, a=100, knn_dist=euclidean, mds_dist=correlation, mds_solver=smacof). (2) 2D UMAP (scanpy, tl.umap, maxiter=3000, spread=3).

<u>Step 2: identifying axes of cellular change:</u>

To robustly identify and model axes of cellular change, we performed a pseudotime analysis using the Palantir^52^ algorithm. In brief, Palantir projects the input data onto a multidimensional diffusion space and constructs a neighbouring graph. It then iteratively refines the shortest paths from a user-defined starting point to each given datapoint, defining this relative distance as the pseudotime. Lastly, it constructs a Markov chain using the neighbouring graph and inferring directionality by the pseudotime. Palantir defines trajectories as the terminal states of the Markov chain and trajectory probabilities as the probability of a given datapoint to reach each of the terminal states.

We ran Palantir over the cellular landscape of all participants (Palantir, run_diffusion_maps and determine_multiscale_ space, n_components=5, knn=50; Palantir, run_palantir, knn=30, max_iterations=100, scale_components=F, n_jobs=1, use_early_cell_as_start=F). We tested multiple starting point for Palantir by random sampling within the cluster of individuals defined as healthy (no decline or AD pathology, as in *Green et al*^6^) that all converged to the same pseudotime.

<u>Step 3: fitting and plotting dynamics:</u>

Dynamics were computed by regressing the feature values over the pseudotime in a specific trajectory using a generalized additive model (GAM). Features used were participants’ traits, mean program score, proportions of community of coordinated programs (defined bellow), expression of single genes, single gene signature score, or AD pathologies. We then used the fitted model to predict the final dynamics over equidistant pseudotime values. In more details, we spline-fitted a GAM for feature *y* in trajectory *j*:

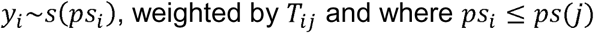

where *i* is a participant, *j* is a trajectory, psi is the pseudotime of participant *i*, *ps*(*j*) is the terminal pseudotime of trajectory *j* and *T_ij_* the trajectory probability matrix of participant *i* in trajectory *j* (mgcv64, gam, formula = *y*∼*s*,(*ps*), weights=*T*_.,*j*_. The final dynamics were predicted over equidistant pseudotime values in the range [0, *ps*(*j*)] (mgcv^53^ predict.gam, se.fit=TRUE). When plotting the dynamics, we presented the predicted feature values over the equidistant pseudotime values, as well as a confidence interval area of predicted value ±2xse, as retrieved from predict.gam.

<u>Step 4: constructing communities of coordinated cellular responses:</u>

To partition the programs into communities with coordinated changes across the aging trajectories, we clustered the programs both by their dynamic patterns and their co-expression across individuals. First, we filtered out the programs that were stably expressed both in AD (prAD) and in the alternative brain aging (ABA) trajectories, to focus on the differential programs. For each program we computed an SNR measurement as the difference between the maximal value of the fit minus the minimal value divided by the maximal standard error (se) over both trajectories.

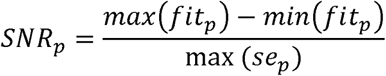

The program inclusion criteria were as follows: if any of the following is true (1) the program is associated with disease trait (amyloid, tangles and cognitive decline) or (2) *sNR_p_* > 10

Partitioning of programs into communities was done by clustering using two similarity measures: (1) similarity in programs dynamics and (2) co-expression of programs across participants. Similarities of dynamics were calculated using a weighted adaptive Radial basis function (RBF) kernel over the z-scored dynamics matrix, computed as follows: We represented each program *p* by its dynamics along both trajectories *x_p_* ∈ ℝ^t1+t2^ for *t*1 and *t*2 the number of equidistant pseudotime values used for the predicted dynamics (see section above). We obtained *x*∼*_p_* by centering and standardizing *x_p_* = (*x_p_* – *mean*(*x_p_*))/*sd*(*x_p_*). We then computed *M* the Mahalanobis distance between every two programs as: *M_pk_* = (*x_p_* – *x_k_*)*^T^W*(*x_p_* – *x_k_*) for *W* a diagonal matrix whose first *t*1 diagonal values are 1/*t*1 and the rest 1/*t*2 (*i.e.* equally weighing both trajectories). Lastly, we calculated the adjacency matrix as *A_dyn_* = *exp*(–*M*/*σσ*^T^), where the division is performed element-wise and clipped to zero values smaller than 10^-4^. *σ* is the vector of local densities at each program calculated as 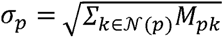 and *N*(*p*) is the set of 5-nearest neighbors of *p* according to *M*. The co-occurrence matrix *A_co_* was calculated as the pairwise spearman correlation matrix of program mean score across all participants (*cor* method, stats package, use=pairwise.complete.obs, method=spearman). That is, [*A_co_*]*_pk_* = *cor* ([*P*]_.*p*_,[*P*]_.*k*_) for filtered version of *P* the abundance matrix defined in the sections above.

We used the Leiden community detection algorithm over the multiplexed 3-layered graph induced by the matrices *A_dyn_*, *A_co_* after zeroing negative correlations and -*A_co_* after zeroing positive correlations (optimise_partition_multiplex method, leidenalg Python package ^54,55^), with layers weighted as 1, 1, −1, respectively. We used the RBERVertexPartition partitioning model^56^, specifying the resolution parameter for each layer as the value maximizing the modularity (resolution_profile method, leidenalg Python package, partition_type=RBERVertexPartition, resolution_range=[10−2, 10], number_iterations=−1). We further refined communities based on the dendrogram calculated within each community based on the dendrogram calculated within each community using *A_dyn_*, *A_co_*.

Once gene expression programs were partitioned into communities of coordinated programs, we assigned each participant a score for every community proportion by averaging the normalized program mean score, and normalized the community proportion vector per participant such that community proportions for every participant sum to one. Community dynamics along trajectories were calculated using the same procedure as used for traits or program dynamics.

#### Dynamics of genes along the BEYOND pseudotime and the inferred trajectories of AD and aging

To fit the dynamics of genes over the gene expression along the pseudotime we computed for each individual the pseudobulk as the average expression of each gene from all cells of this individual. (Seurat AverageExpression, assays=’SCT’, group.by=’projid’). Then each gene’s average expression was converted to z-scores across individuals and fitted the dynamics fitted by GAM as described above.

For visualization of dynamics of multiple genes we converted the predicted values per gene to z-scores. We also computed the log fold-change, computed as the log_2_ of the maximal value of the fit divided by the minimal value of the fit (plus epsilon) over the pseudotime of the trajectory:

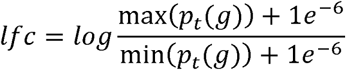

were *p_t_* (*g*) is the z-scored value of the predicted value of the gene *g* at the pseudotime *t*.

To fit the dynamics of a specific pathway along the pseudotime - we first defined the gene set as genes within the pathway that were differentially expressed in the relevant program signatures, specifically for the corresponding AD-associated programs we included genes in the signatures of the three neuronal classes within the relevant program. Next, we applied z-score transformation for each gene expression levels over the pseudobulk of individuals (as defined above), and defined the pathway expression per individual as the mean of the scaled values over all pathway genes. Finally, we fitted the dynamics of the pathway expression across individuals by GAM (as described above).

### Comparison of cell-cycle reentry and the stress-apoptosis decreasing neuronal programs

#### Quantification of neuronal cells expressing disease-programs

To quantify the number of neurons that express one of the disease-programs (*i.e.* the *Stress/Apoptosis* or *Cell-cycle reentry* programs) we defined two sets of individuals: a control set - healthy group of individual with no/low AD-pathologies and cognitively healthy defined by their assigned position at the beginning of the ageing pseudotime (pseudotime < 0.25), and an AD set - diseased group of individuals with abundant AD-pathologies and cognitively declined defined by their assignment at the end of the prAD trajectory (pseudotime > 0.6 & prAD probability > 0.9). Since the scoring of the expression programs within each cell is a continuous value, to define which cells are positively expressing a program, we defined a threshold for expressing it using the top quantile over the distribution of the program scores in all neuronal cells from the control se of healthy individuals (not expected to express the disease programs). All neurons, within the control and the AD set, with scores above this threshold were annotated as expressing the disease program, and termed *Stress/Apoptosis* or *Cell-cycle reentry* cells for the respective program. We carried the analyses for different quantiles, 95%, 90% and 85%, and showed consisted conclusions for all these thresholds. We next computed the percent of cells expressing each of the two programs out of the total number of neurons per individual, and out of the total number of neurons within a neuronal class or a neuronal subtype.

#### Scoring synaptic-gene expression across groups

To compare the signature score of synaptic genes we took all the differentially expressed genes from the pathways GO:0098693, hsa04721, and GO:0099504 (regulation of synaptic vesicle cycle, Synaptic vesicle cycle and synaptic vesicle cycle) for presynaptic signature and GO:0099572 (postsynaptic specialization) for postsynaptic genes.

Using Seurat’s AddModuleScore function over the inhibitory, excitatory L2-3 and excitatory L4-6 we scored each sets of genes. Of note, when comparing the neurotransmitters expression, we compared their expression only within the relevant neuronal subtype, *i.e.* SST and VIP neurons for the genes SST and VIP respectively.

We used Wilcoxon-test (wilcox.test) to test the difference in the expression between groups of individuals/cells – comparing within the control set and the AD set the *Cell-cycle reentry* expressing-neurons vs. *Stress/Apoptosis* expressing neurons (defined as above). We applied Spearman correlation (stats, cor.test, method=spearman) to test the correlation between neuronal programs scores and gene expression or gene pathway score. We then applied FDR multiple hypothesis correction (p.adjust, methods=BH) and set the alpha for 0.05.

## Supporting information

Supplementary Table 1 - Cohorts description

Supplementary Table 2 - Programs characterization

Supplementary Table 3 - Abundance and Endophenotypes Associations

Supplementary Table 4 - BEYOND analysis results

Supplementary Table 5 - AD-associated programs

## Data availability

SnRNA-seq data are available via the AD Knowledge Portal (https://adknowledgeportal.org), in the Synapse database: https://www.synapse.org/#!Synapse:syn31512863

Proteomics and other ROSMAP resources can be requested at the RADC Resource Sharing Hub at: https://www.radc.rush.edu. Source data are provided with this paper.

## Code availability

The code used in this study to model gene expression programs, accompanying analysis and generation of all figures is available at: https://github.com/naomihabiblab/continuous-modeling

## Acknowledgements

We thank the individuals who donated their brain to research through the Rush University Alzheimer’s Disease Center. The work was supported by NIH RF1 AG057473 (P.L.D.J. and D.A.B.), U01 AG061356 (P.L.D.J. and D.A.B.), U01 AG046152 (P.L.D.J. and D.A.B.), R01 AG070438 (P.L.D.J.), P30AG10161, P30AG72975, R01AG17917, R01 AG015819 (D.A.B.), U01 AG072572 (P.L.D.J.). The Israel Science Foundation (ISF) research grant no. 1709/19, the European Research Council grant 853409, the and the Myers Foundation (N.H.); G.S. was supported by the JBC fellowship. Diagrams in Figs 1a and 2c were created using BioRender.

## Contributions

N.H., R.M. and G.S.G. designed the study. R.M. and G.S. performed the computational and statistical analyses, with guidance of N.H. and with the help of G.S.G, A.A.L. M.A. and A.C. and with input from V.M. and P.L.D.J. N.H. R.M. and G.S. wrote the manuscript, and all of the co-authors provided critical comments. D.A.B. is principal investigator of the parent ROS and MAP studies and performed study supervision.

## Notes

### Competing Interest Statement

The authors have declared no competing interest.

