## Supplementary Figures for "Early neuronal reprogramming and cell cycle reentry shape Alzheimer’s disease progression"

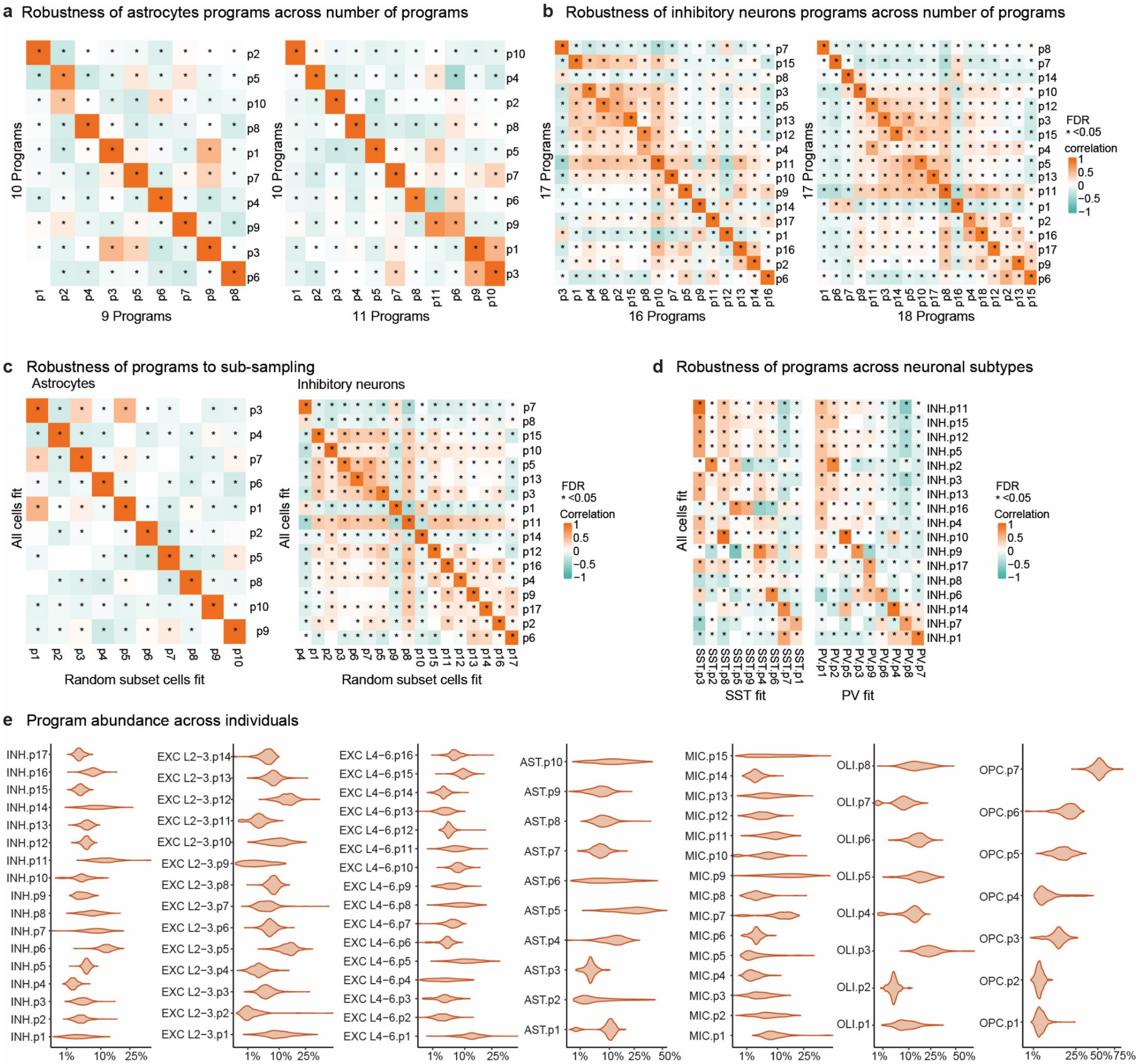


**Extended Data Fig 1. Robustness and distribution of co-expression programs.**

**a,b.** Robustness of programs over selection of number of programs (k). Spearman correlation of the programs scores across cells for different number of learned programs, shown for astrocytes and inhibitory neurons (FDR corrected p-value). **c.** Robustness of programs over random subsets of cells within a cell-type. Spearman correlation of program scores across cells when fitting the programs on a random subset of 10,000 cells compared to all the cells, shown for astrocytes and inhibitory neurons (FDR corrected p-value). **d**. Robustness of shared neuronal programs. Spearman correlation of program scores across cells when fitting programs across all inhibitory neurons or separately for specific inhibitory neuronal subtypes, shown for SST and PVALB subtypes (FDR corrected p-values). **e.** Programs abundance across individuals for inhibitory neurons, excitatory lower-layer neurons, excitatory upper layers neurons, astrocytes, microglia, oligodendrocytes and OPCs.

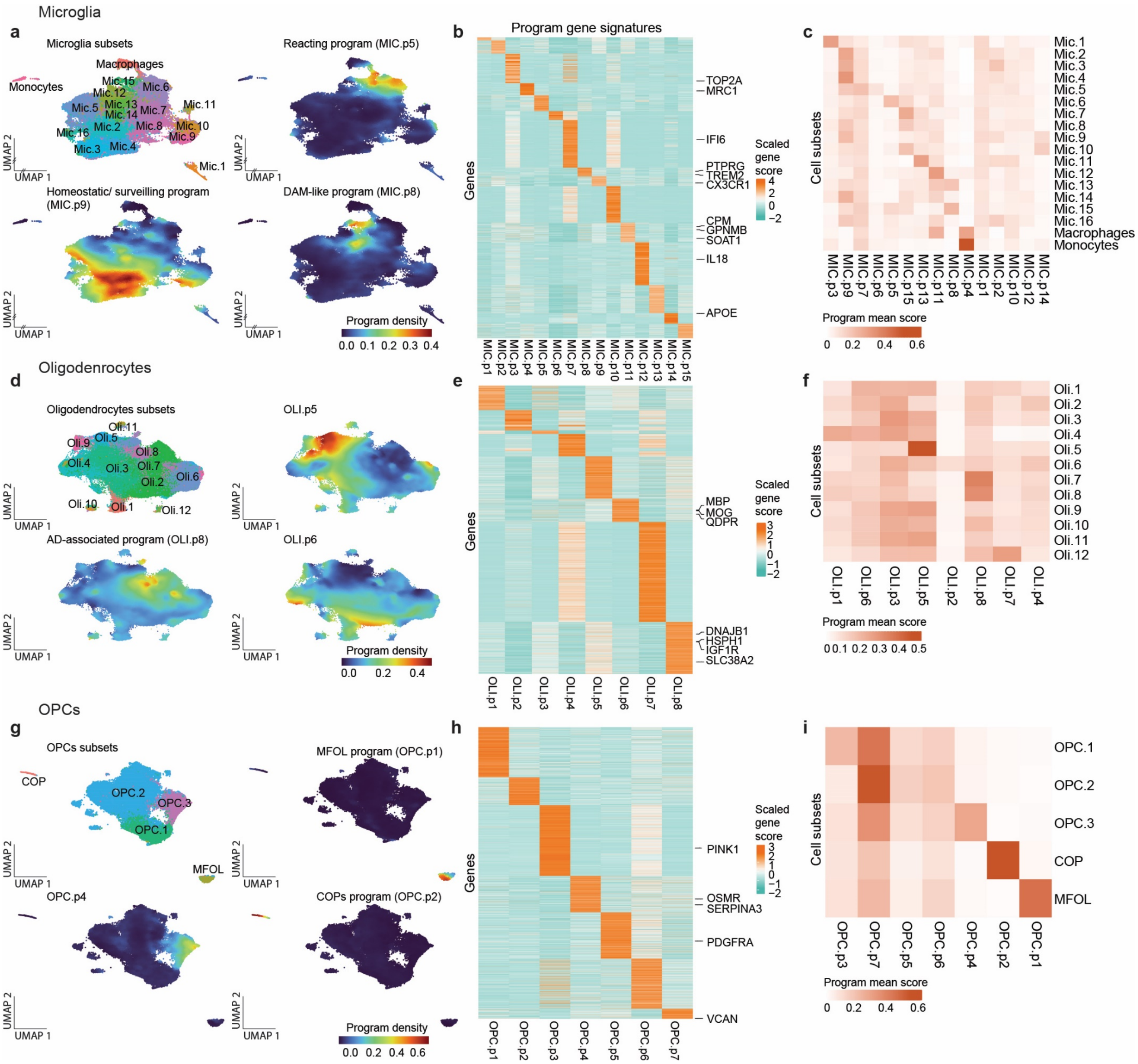


**Extended Data Fig 2. Alignment of cell-types specific programs with cluster annotations.** **a,d,g.** 2d UMAP embedding of snRNA-seq profiles coloured by *Green et al.*[6] subsets (clusters, up-left) and by locally smoothed density values of representative programs for microglia (**a**, n=86,612)**,** OPCs (**d**, n=62,979) and oligodendrocytes (**g**, n=346,593). **b,e,h.** Unique expression signatures across programs. Gene score (F, **Methods**) for top differentially expressed genes (rows, scaled) across programs for microglia (**b**)**,** OPCs (**e**) and oligodendrocytes (**h**). **c,f,i.** Programs alignment with previously defined subsets (clusters). Averaged program scores within each cell subset from *Green et al*[6] for microglia (**c**)**,** OPCs (**f**) and oligodendrocytes (**i**).

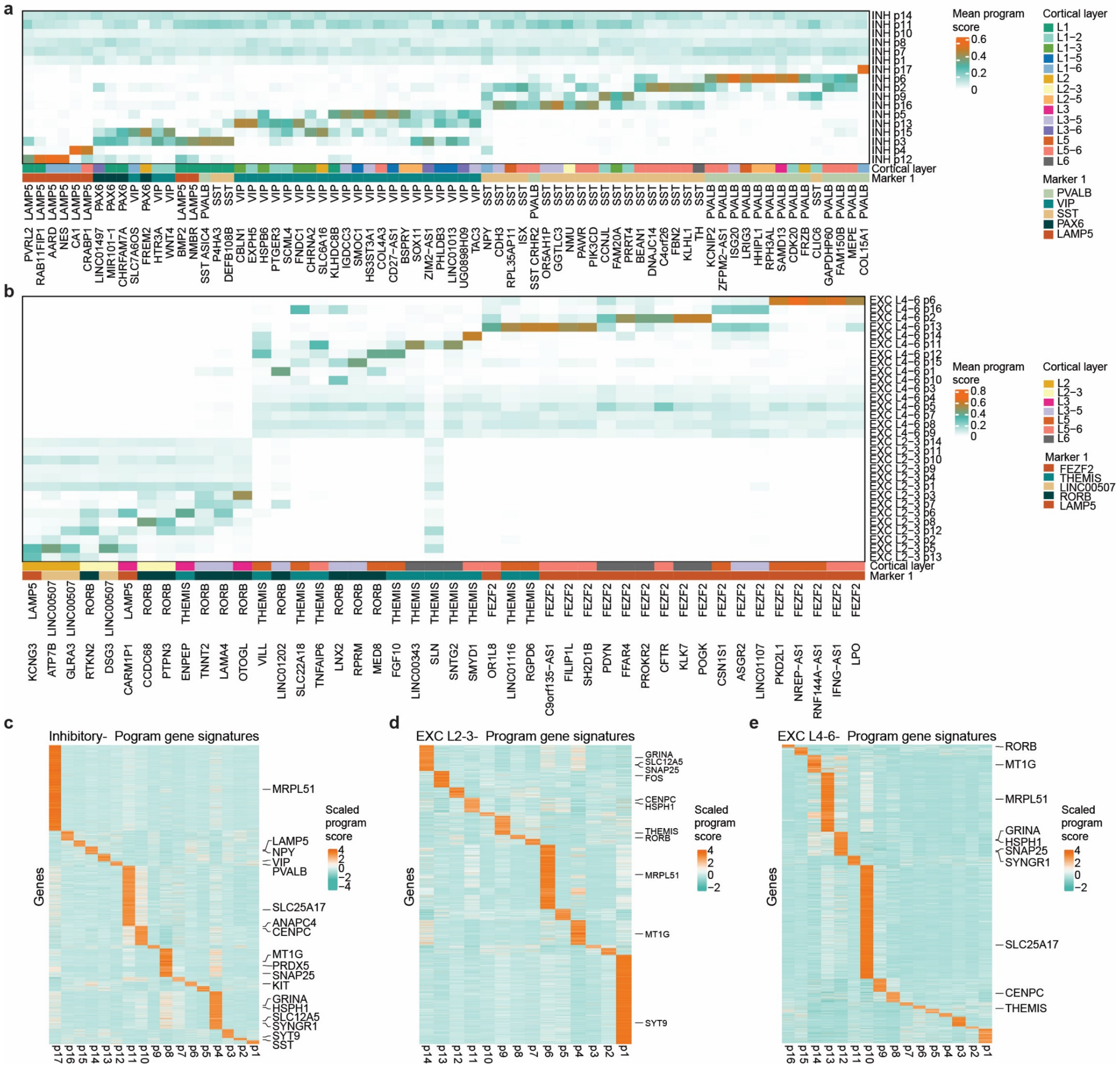


**Extended Data Fig 3. Alignment of neuronal programs with subtype annotations.**
**a,b.** Association of inhibitory (**a**) and excitatory (**b**) neuronal programs to Allen Brain Map neuronal annotations[10]. Averaged program scores within each annotated neuronal subtype, highlighting subtypes-specific programs and shared programs across subtypes. Colour bars indicate cortical layers and the markers of each neuronal subtype. **c-f.** Unique expression signatures across programs. Gene score (F, **Methods**) for top differentially expressed genes (rows, scaled) across programs for inhibitory (**c**) excitatory lower layers (**d**) and excitatory lower layers (**e**) neurons.

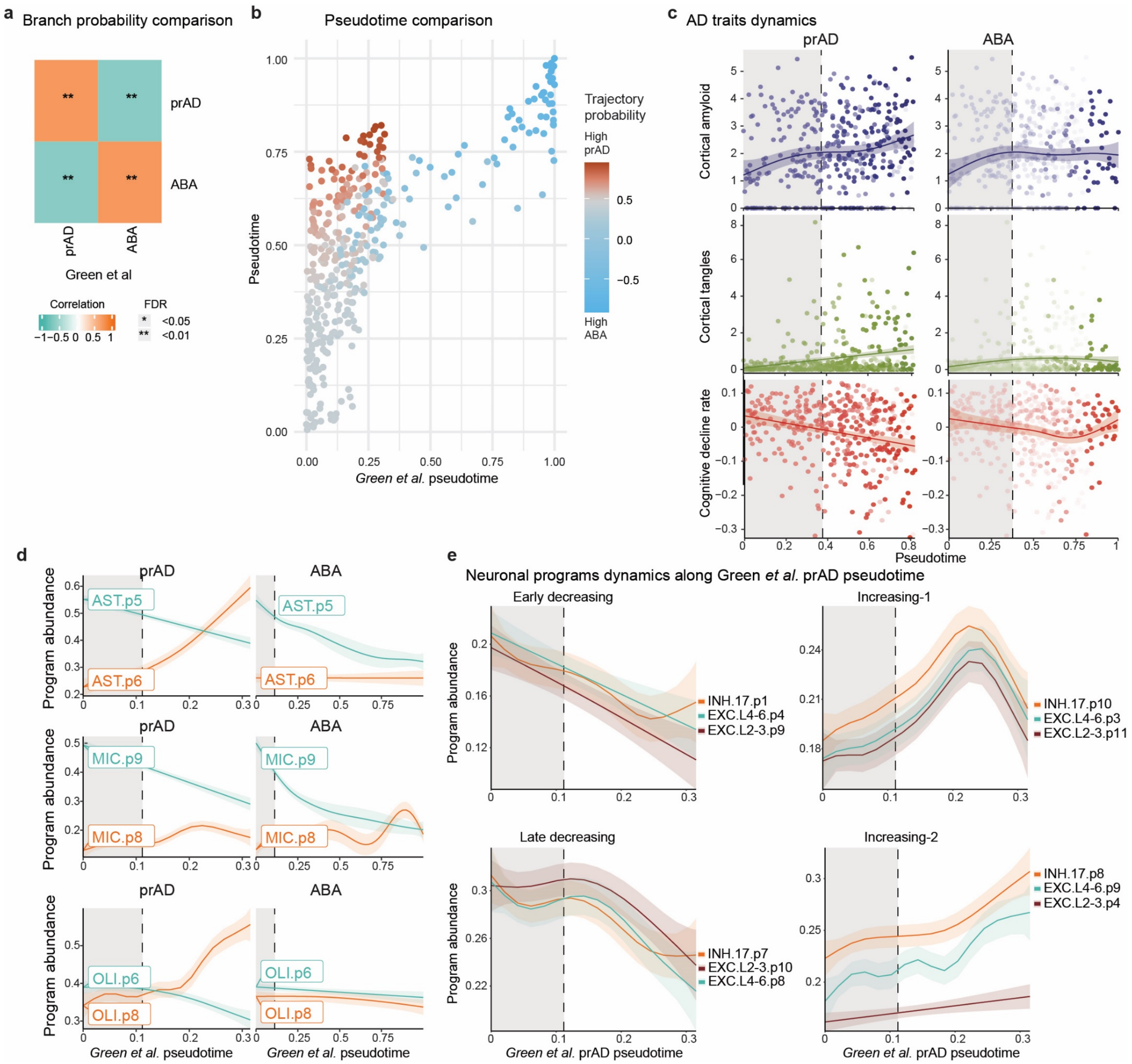


**Extended Data Fig 4. Program-based trajectories align with cluster-based inference.**

**a,b.** Comparison of BEYOND trajectories inferred over programs and over clusters (from *Green et al.*[6]). **a.** Correlation of trajectory assignment probabilities of individuals **b.** Comparison of pseudotime assignments for each individual, coloured by the probability of being in the prAD trajectory. **c.** Dynamics of AD traits along the prAD and ABA pseudotime (as in **Fig. 2d**). Lines are the inferred dynamics of the traits per individual. Dots are individuals coloured by the probability per trajectory. Dashed line represents the split point between trajectories. (**d-e**) Dynamically regulated glial and neuronal programs calculated over *Green et al* pseudotime (compared to **Fig. 2e-f**). Inferred dynamics of averaged program scores per individual over the pseudotime. Dashed line represents the split point between trajectories. (**d**) Decreasing homeostatic and increasing disease-associated programs in astrocytes (top), microglia (middle) and oligodendrocytes (bottom). (**e**) Corresponding neuronal programs across classes: early-decreasing (top left), late-decreasing (bottom left), increasing-1 (top right) and increasing-2 (bottom right).

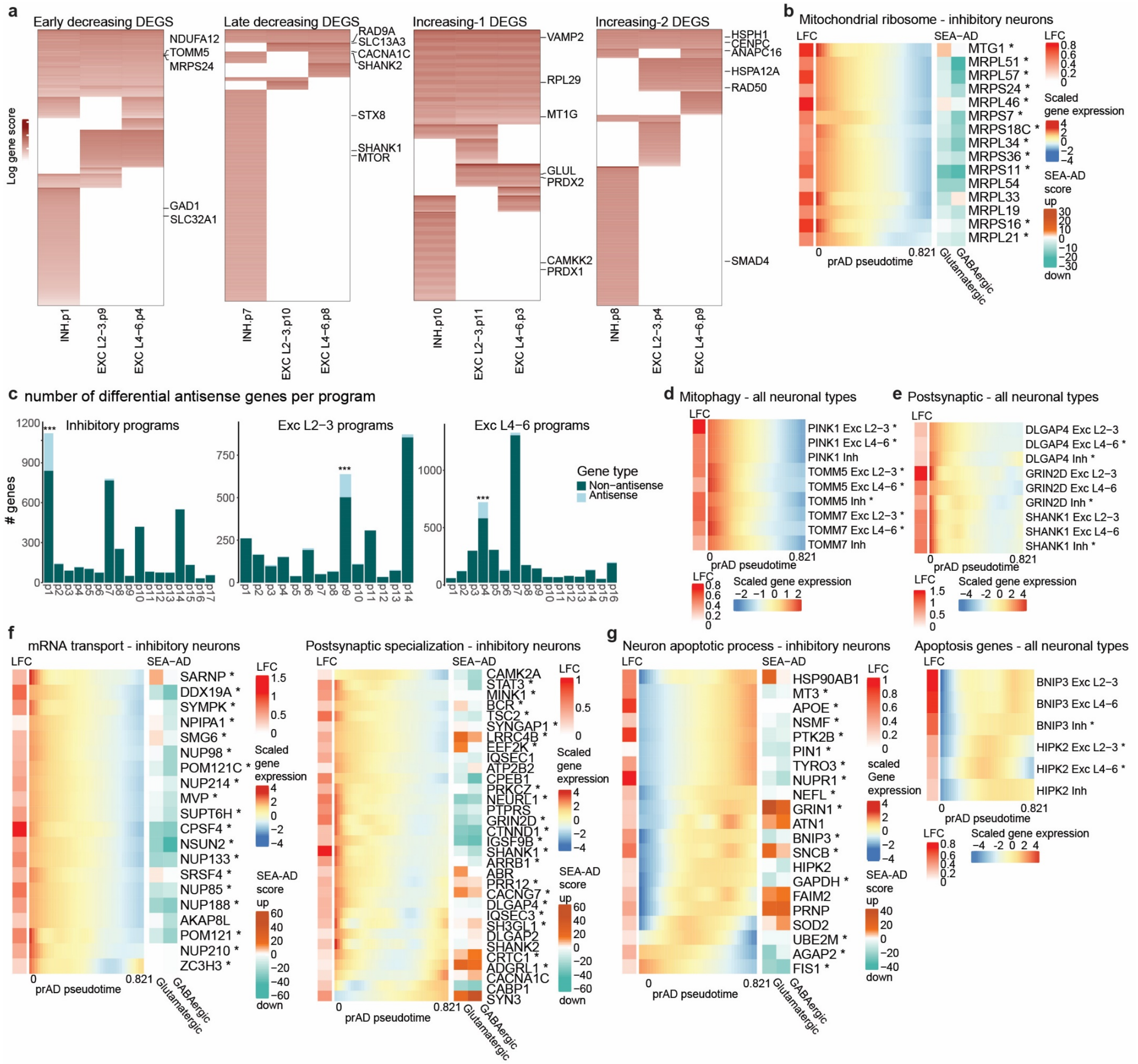

**Extended Data Fig 5. Annotation and validation of AD-associated neuronal programs.**
**a.** Shared and unique genes within each corresponding group of neuronal programs between inhibitory, excitatory L2-3, and excitatory L4-6 neurons. **b.** Mitochondrial-ribosome genes (enriched in INH.P1) dynamics and validation in an independent snRNA-seq atlas. Left: log fold change (LFC) of the gene expression along pseudotime (**Methods**). Middle: Gene expression dynamics along the prAD trajectory in inhibitory neurons. Coloured by the inferred expression level along the pseudotime, scaled per gene. Asterisk denote genes differentially expressed in the program. Right: Score over number of neuronal subtypes that differentially expressed the noted gene in SEA-AD data[7], calculated as the difference between the number of subtypes that upregulated the gene and the ones that downregulated it (**Methods**). **c.** Enrichment for antisense genes in inhibitory, excitatory L2-3 and excitatory L4-6 programs (hyper-geometric test, FDR<0.001, **Methods**). **d,e.** Mitophagy (**d**) and postsynaptic (**e**) gene dynamics along the prAD trajectory across neuronal classes. Presented as in b. Asterisk denote differentially expressed genes within the program. **f.** Dynamics and validation in an independent snRNA-seq atlas[7] of pathways linked to the late-decreasing inhibitory program INH.p7. Showing mRNA transport and

postsynaptic specialization genes as in b. **g.** Dynamics and validation in an independent snRNA-seq atlas[7] of pathways linked to the increasing inhibitory program INH.p10. Left: Neuron apoptotic process shown as in b. Right: Dynamics of key apoptotic genes across all neuronal types shown as in d.

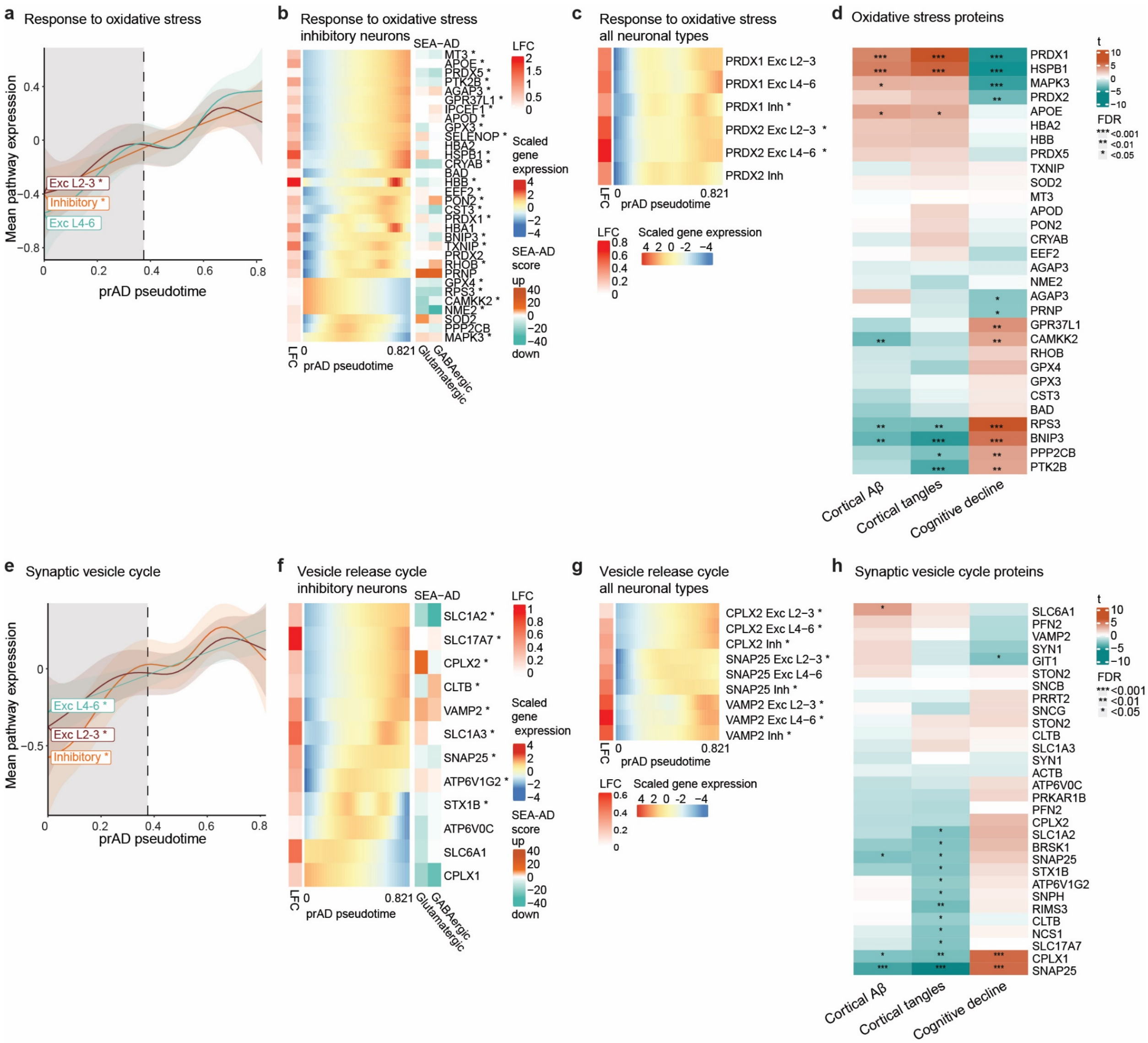

**Extended Data Fig 6. Annotation and validation of AD-increasing neuronal programs.**
**a-d.** Validation and dynamics of response to oxidative stress genes enriched in INH.P10. **e-h.** Validation and dynamics of genes from synaptic vesicle pathways enriched in INH.P10. **a,e.** Inferred dynamics of pathway signature genes along the prAD trajectory across neuronal classes. Inferred dynamics over the pseudotime of averaged scaled pathway expression (**Methods**) across cells per individual. Dashed line represents the split point between trajectories. Asterisk denotes pathway significantly enriched in this program. **b,f.** Dynamics and validation in an independent snRNA-seq atlas of the genes from the enriched pathway. Left: log fold change (LFC) of the gene expression along pseudotime (**Methods**). Middle: Gene expression dynamics along the prAD trajectory in inhibitory neurons. Coloured by the inferred expression level along the pseudotime, scaled per gene. Asterisk denotes genes differentially expressed in the program. Right: Score over the number of neuronal subtypes that differentially expressed the noted gene in SEA-AD data[7]. **c,g.** Gene dynamics along the prAD trajectory across neuronal classes. Presented as in b. Asterisk denotes differentially expressed genes within the program. **d,h**. Validation of neuronal expression changes in AD by proteomics. Association between proteins and AD traits of the pathway genes (n=400, FDR corrected, **Methods**).

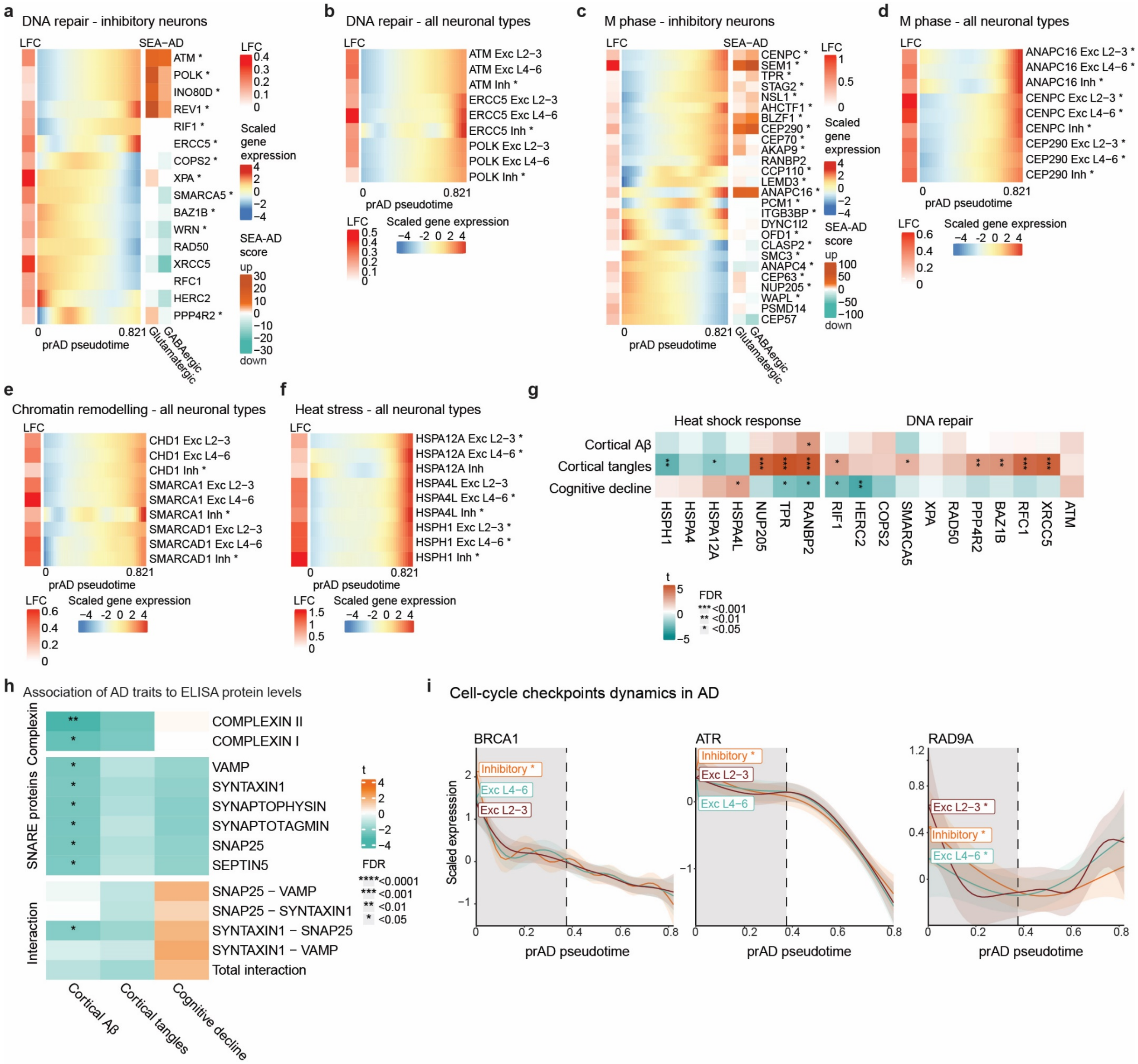


**Extended Data Fig 7. Dynamics and validation of AD-associated neuronal programs and cell-cycle reentry checkpoints.**

**a,b.** Dynamics and validation in an independent snRNA-seq atlas[7] of DNA repair genes (a) and M-phase genes (b) linked to the increasing inhibitory program INH.p8. left heatmap: Left: log fold change (LFC) of the gene expression along pseudotime (**Methods**). Middle: Gene expression dynamics along the prAD trajectory in inhibitory neurons. Coloured by the inferred expression level along the pseudotime, scaled per gene. Asterisk denotes genes differentially expressed in the program. Right: Score over the number of neuronal subtypes that differentially expressed the gene in SEA-AD data[7]. Right heatmap: Scaled gene expression dynamics along the prAD trajectory across neuronal classes. **c,d**. Dynamics of chromatin remodelling (**c**) and heat stress (**d**) genes linked to the increasing cell-cycle reentry programs across neuronal classes. Presented as in (**a**). **e**. Proteomics-based validation of neuronal expression changes of heat-shock and DNA-repair genes in AD. Association between proteins and AD traits of the pathway genes (n=400, FDR corrected, **Methods**). **f**. Association between ELISA synaptic proteins and pairs of interacting proteins levels and AD traits (n=192, FDR corrected, **Methods**). **g**. Inferred dynamics of cell-cycle checkpoints genes across neuronal classes. Asterisk denotes if the gene is differentially expressed in the cell-cycle reentry program within each neuronal class.

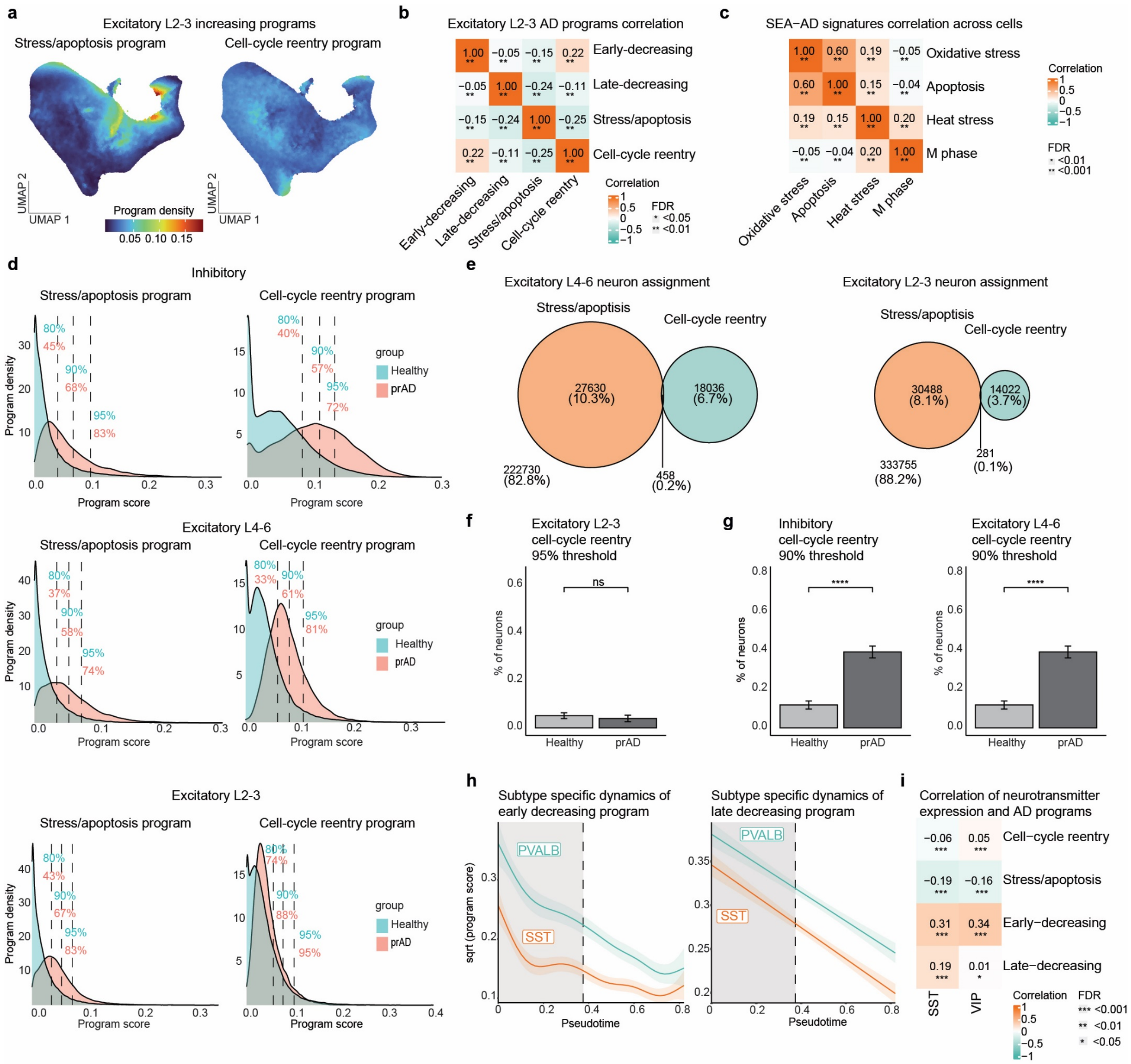


**Extended Data Fig 8. Mutual exclusive expression of the cell-cycle reentry and the stress/apoptosis neuronal programs linked to neuronal vulnerability (accompanying Fig. 4).**

(**a-c**). Mutually exclusive expression of the cell-cycle reentry and the stress/apoptosis neuronal programs. **a.** UMAP embedding of excitatory L2-3 neurons, coloured by locally smoothed density value of each program per cell. **b**. Correlations of the decreasing and increasing disease-associated programs across excitatory L2-3 neurons. Spearman correlations with FDR-adjusted significance. **c.** Spearman correlation of signature scores of key pathways in the cell-cycle reentry and stress/apoptosis programs in SEA-AD data[7] (**Methods**) **d.** Distribution of program scores across cells from healthy compared to advanced AD individuals. Distribution of stress/apoptosis (left) and cell-cycle reentry (right) scores within inhibitory (top) excitatory L4-6 (middle) and L2-3 (bottom) neurons in healthy and advanced AD groups of individuals (**Methods**). Dashed lines indicate different quantiles of program score, coloured advanced AD or healthy. **e.** Quantification of excitatory L4-6 and L2-3 neuronal cells assignment to stress/apoptosis or cell cycle reentry showing minimal overlap (threshold of 95%, **Methods**). **f-g.** Average proportions of cell-cycle reentry neurons in healthy (n=78) and advanced prAD (n=34) groups of individuals (**Methods**).

Wilcoxon test p-value<0.001. Showing excitatory L2-3 with 95% quantile (**f**, along with **Fig. 4e**), and inhibitory and excitatory L4-6 with 90% quantile (**g**). **h.** Dynamics of early-decreasing (left) and late-decreasing (right) average program scores along the prAD trajectory for relative vulnerable SST and resilient PVALB inhibitory neuronal sub-types. **i.** Spearman correlation of neurotransmitters expression in related inhibitory neuronal subtypes (SST n=55563, VIP n=61586) to AD-associated program scores (FDR-adjusted).

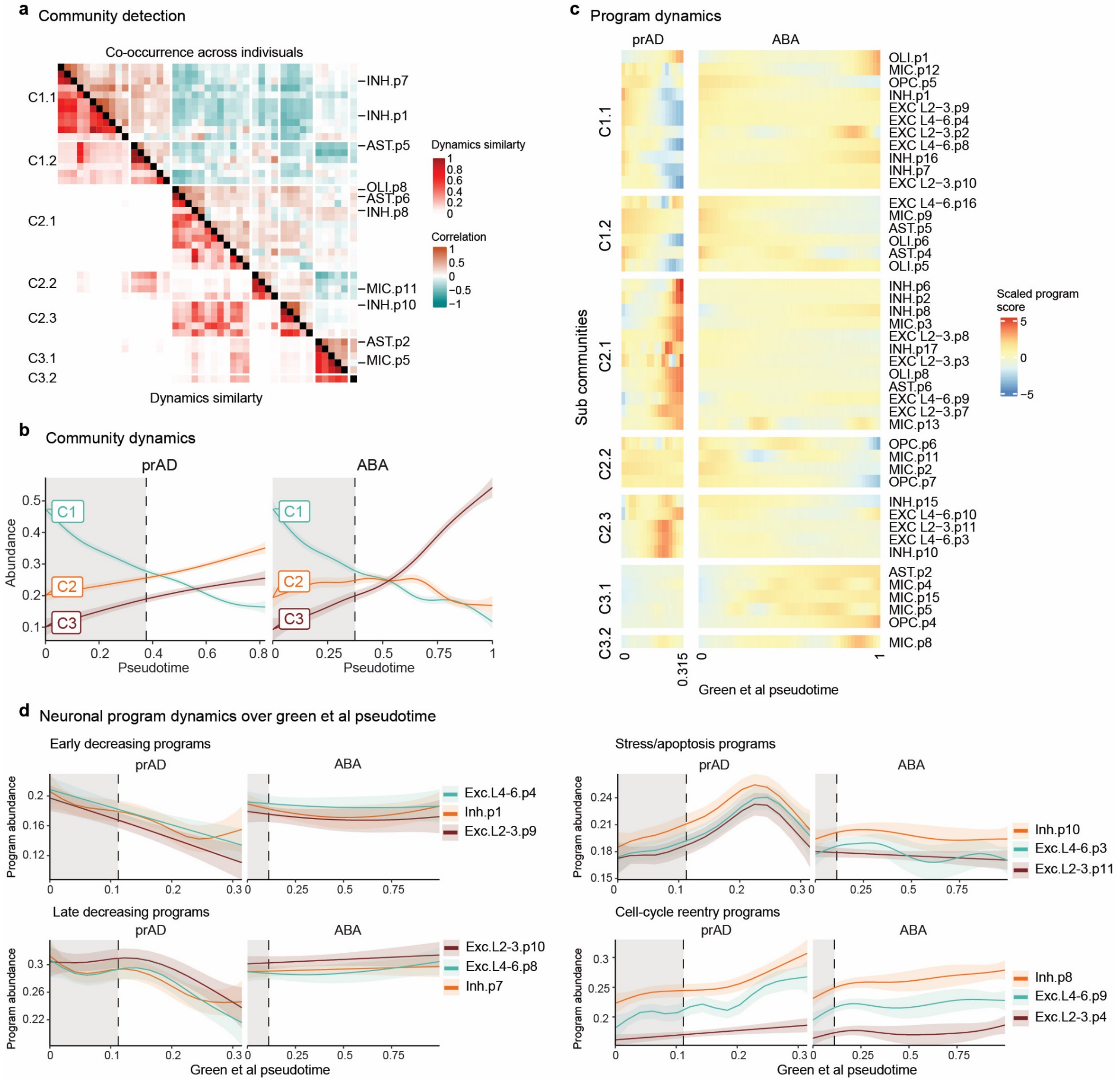


**Extended Data Fig 9. Distinct modulation of coordinated neuronal and glial community of programs in AD.**

**a.** Programs similarity measures: pairwise correlations of program abundance (co-occurrence, top) and dynamics similarity (bottom, **Methods**). Programs are grouped by sub-communities (**Methods**). **b.** Distinct dynamic patterns of communities along the two trajectories (prAD, left; ABA, right). Inferred dynamics over averaged of programs scores within the community (**Methods**). Dashed line represents the split point between trajectories. **c.** Inferred program dynamics along the published trajectories in *Green et al.*[6]. Program dynamics along the prAD pseudotime (left) and the ABA pseudotime (right) divided into 7 sub-communities (as in **Fig. 4a**, **Methods**). Colour by the inferred program abundance scaled along the pseudotime. **d.** AD-associated neuronal programs are specifically modulated in the prAD trajectory and not in the ABA along the published trajectories in *Green et al.*[6]. Dynamics of the groups of corresponding decreasing and increasing neuronal programs along the prAD trajectory (left) and the ABA trajectory (right) of *Green et al.* [6].
